# Disentangling urbanisation, climate effects and their interaction on ornamental colourations

**DOI:** 10.1101/2025.02.19.639098

**Authors:** Lisa Sandmeyer, David López-Idiáquez, Amélie Fargevieille, Pablo Giovannini, Samuel Perret, Maria del Rey, Anne Charmantier, Claire Doutrelant, Arnaud Grégoire

## Abstract

Urbanisation and climate change can both affect phenotype expression across taxa. Most evidence collected so far has focused on exploring these two phenomena on isolation. Currently, the combined effects of climate change and urbanisation remain underexplored, despite being among the greatest challenges faced by biodiversity. Here, we use a decade-long, individual-based study of urban and forest great tits (*Parus major*) to analyse urbanisation, climate and their interactive effects on yellow breast colouration, a carotenoid-based trait. We find that urban birds exhibit duller colourations than forest counterparts, being 10-20% less chromatic, with first-year birds and males being more negatively impacted by cities. Additionally, birds in the city are more sexually dichromatic than in the forest. Over the decade, colouration differences between habitats remain stable, following a similar quadratic temporal pattern. Finally, while climate has a weak effect on colouration, urban birds appear more sensitive to its influence than forest birds. Our results indicate urbanisation has a stronger impact than last decade climatic variation on great tit colouration, though both factors may interact. The heightened sensitivity of first-year birds and increased sexual dichromatism in cities may alter the strength of natural and sexual selection on this trait in urban environments.

## Introduction

Urbanisation and climate change are two major factors driving ongoing environmental changes in wild populations worldwide (Alberti et al., 2017; Thackeray et al., 2016). Research across several taxa shows both plastic and genetic responses to these anthropogenic changes (e.g. Lambert et al. 2021; Radchuk et al. 2019). The relevance of urbanisation as an environmental driver of phenotypic change has increased in recent decades with predictions that by 2030, urban land cover will triple compared to its size in 2000 (Seto et al., 2012). Urban landscapes are associated with large environmental changes such as increased pollution (chemical: Isaksson 2010; acoustic: Slabbekoorn and Peet 2003; light: Longcore and Rich 2004) and changes in food availability and/or quality (Szulkin et al., 2020). These novel environmental conditions are expected to impose novel selective forces that can shape phenotypic variation of populations living in urban environments (Luder et al., 2018; Seress et al., 2018; Thompson et al., 2022). Phenotypic divergence in animals has been reported in traits such as reproductive phenology, morphology, and behaviour, with urban populations usually being earlier breeders, smaller and bolder than their rural counterparts (Capilla-Lasheras et al., 2022; Eggenberger et al., 2019; Thompson et al., 2022). Long-term evolutionary ecology research has also revealed the importance of climate change as an environmental driver of phenotypic changes (Parmesan and Yohe 2003). For instance, rapid warming is causing advancements in reproductive timing and reduction in body size across different taxa (Hällfors et al., 2020; Kellermann & Van Heerwaarden, 2019; Thackeray et al., 2016).

One of the most striking environmental changes in cities, when compared to natural habitats, is the urban heat island effect. This phenomenon is characterized by cities consistently experiencing higher temperatures than their surrounding rural areas (Oke, 1982). The urban heat island effect has been linked to the evolution of thermal tolerance in species (*e.g.,* Diamond et al. 2018), and to changes in breeding phenology in several taxa (Li et al., 2019; Merckx et al., 2021). Urbanisation effects on ambient temperature can be similar in magnitude to those of climate warming, but they lead to spatial rather than temporal variation in temperature. As such, urban areas can serve as models for studying the consequences of climate change (Verheyen et al., 2019). Additionally, climate change effects may be more pronounced in urban populations than in their rural counterparts due to interactive effects with the urban heat island and other urban stressors (Urban et al., 2024).

A small number of studies have investigated the interaction between climate change and urbanisation. Results showed exacerbated effects of urbanisation under high temperatures and climate change on the phenology of several species of butterflies as well as buffering effects on the reproduction of great tits (Pipoly et al. 2022 for reproduction; Diamond et al. 2014 for phenology; but see Sumasgutner et al. 2023). For example, Pipoly et al. (2022) reported a reduction in great tit reproductive success as a consequences of extreme heat events in forests but not in cities. However, besides phenology or body condition, other phenotypic traits can also be sensitive to the environmental changes induced by urbanisation and climate change. Condition-dependent signals, such as ornaments, are an example of this. Due to their tight link to individual condition and quality (Dougherty 2021) they are expected to be impacted by both climatic and urban effects, still this remains to be evaluated.

Ornaments, such as conspicuous colourations or complex songs, play a crucial role shaping interactions between individuals in inter and intra-sexual contexts (Andersson, 1994; Maynard Smith & Harper, 2003). Their expression often relies on the production and maintenance costs linked to the trait (Cotton et al., 2004) and consequently, ornaments are expected to be associated with variation in environmental conditions (Hill, 1995). In birds, for instance, carotenoid-based colours (i.e., those generated by depositing carotenoids on feathers) are strongly tied to environmental factors, as birds cannot synthetize carotenoids *de novo* and must acquire them through diet (Goodwin, 1984). Carotenoid-based colourations are widespread in birds; approximately 40% of passerine species deposit carotenoids on their feathers and around 50% of non-passerine also show carotenoid-based colours in other integuments (Davis & Clarke, 2022; Thomas et al., 2014). Carotenoid colouration may convey information about an individual’s ability to reproduce, to secure a territory or its health status (Hill et al., 2023; Weaver et al., 2018). For example, carotenoid-based colouration has been linked to parasite prevalence (Badiane et al., 2022; Del Cerro et al., 2010), rearing capacity (Brown et al., 2014; García-Campa et al., 2022), resistance to oxidative stress (Cote et al. 2010, Pérez-Rodríguez, Mougeot, and Alonso-Alvarez 2010) and temperature (Breckels & Neff, 2013; Eraud et al., 2007). It can also be involved in secondary sexual signalling, as a cue of quality for potential mates (intersexual selection) or of ability to acquire and defend a breeding territory (intrasexual selection) (Svensson & Wong, 2011). Because of this high environmental sensitivity and key functions, it has been suggested that they may be useful as bio-monitors (Koneru & Caro, 2022; Peneaux, Grainger, et al., 2021). Understanding impacts of urbanisation and climate on such ornaments could thus help in future conservation or environmental impact studies.

Research on urban effects on carotenoid-based colouration has described an “*urban dullness*” phenomenon, whereby urban avian populations tend to show duller carotenoid-based colourations than their rural counterparts (Leveau, 2021). For example, great tits (*Parus major*) are known to be duller in European urban compared to forest populations, but there is variation in this effect depending on the urban-forest pair investigated (Biard et al., 2017; Isaksson et al., 2005; Salmón et al., 2023). Two main direct environmental drivers of these urban colour shifts have been discussed in the literature: food and pollution, with food potentially being indirectly affected by temperature, rainfall, and forest cover. First, food availability and quality are crucially different across the two habitats. Caterpillars living in urban settings contain lower levels of carotenoids and are less numerous than their forest-dwelling counterparts (Isaksson & Andersson, 2007; Pollock et al., 2017). Moreover, although anthropogenic food sources can be abundant in cities (Coogan et al., 2018), their nutritional composition may be ill-fitted for urban individuals (Pollock et al., 2017). Second, increased stress in cities resulting from different sources of pollution (*e.g.,* chemical, light) could increase the use of carotenoids for keeping an optimal physiological status for urban compared to rural individuals, reducing carotenoid availability for colouration in city birds (Giraudeau et al., 2015 in house finches *Haemorhous mexicanus*; Isaksson, 2015).

Higher temperatures, due to climate and urbanisation, could increase the relative costs associated with maintaining enhanced carotenoid-based colourations through three main pathways. First, higher temperatures can reduce plant carotenoid concentration (Dhami & Cazzonelli, 2020), thus reducing carotenoid availability. Second, higher temperatures increase chemical pollution, which may increase physiological stress in plants and birds (Viatte et al., 2022). Finally, higher temperatures induce a physiological challenge depending on the animal’s thermal tolerance, especially during extreme climatic events, with potential impacts on an individual’s colouration (e.g., Diamond et al., 2018; Sumasgutner et al., 2023). Effects of climate change on carotenoid-based colours are still rarely investigated, and the studies to date provide contrasting results. For instance, in a 15-year study on a population of blue tits (*Cyanistes caeruleus)* from southern France (Corsica), increased temperatures were associated with a decrease in yellow colouration (López-Idiáquez et al., 2022), while it was linked to an increase in the yellow colouration of a great tit population from central Europe (8-year-study in Hungary, Laczi et al., 2020). Although this suggests that bird carotenoid colouration is sensitive to temperature variation, it also shows that climate change effects can differ among populations. In particular, Southern Europe populations might be more sensitive to climate change, since heat pressures in this region can be higher than those experienced at higher latitudes (Giorgi, 2006). To our knowledge, no study has simultaneously examined variations in urbanisation and climate to understand their potential additive (*i.e.,* synergetic) or interactive (*e.g.,* antagonistic) impact on colouration.

Here we addressed these questions using 10 years of data including 1606 measurements (for 604 females and 567 males) of great tit carotenoid-based colouration in a pair of urban and forest populations situated in southern France. We focused on three colouration measures: yellow chroma (or colour intensity) linked to the quantity of carotenoids deposited on the yellow feathers (Isaksson et al., 2008), and mean brightness and UV chroma, both linked to the structure of the yellow feathers (Prum, 2006). Firstly, we tested whether forest great tits differed in these three coloured parameters compared to their urban counterparts. Secondly, we analysed temporal trends of great tit colouration across our study period to investigate whether colouration has showed a change that could be aligned with climate change. Lastly, we analysed temporal changes in temperature and precipitation over the same period and explored the links between colour, climate, and urbanisation. Overall, we predicted that urban great tits will show duller breast colourations than those living in the forest. We also predicted a temporal trend in colouration along our study period, with a decrease in colouration, as a response to an expected increase in temperature and decrease in precipitations (López-Idiáquez et al., 2022). Since we expected a differential sensitivity to climate in urban areas, we predicted more pronounced effects of temperature in the urban habitat than in the forest.

## Material and methods

### Study areas and general methods

The study was conducted between 2013 and 2022 in two great tit populations in southern France. The first study area is the city of Montpellier (43°36’55” N, 3°52’16” O, hereafter city, Figure S1) and includes eight sites capturing different degrees of urbanisation (*i.e.,*vegetation cover, car traffic, pedestrian presence and artificial light, urban parks and trees on roadsides, for details: Caizergues et al., 2021; Demeyrier et al., 2016). The second, is located in an oak forest in Montarnaud (La Rouvière, 43°59’53” N, 3°40’9” O, hereafter forest, Figure S1), approximately 20km northwest from Montpellier. We captured breeding great tit pairs when nestlings were 10 to 15 days old, and ringed them with a uniquely numbered metal ring. At that time, we aged each parent as yearlings (one year old) or adults (≥2 years old) based on their plumage (Svensson, 1992). We sexed the birds using a combination of plumage features and the presence of a brood patch. To measure colouration, we plucked eight feathers from the upper part of their yellow breast patch following a standardised protocol (Doutrelant et al., 2008). The feathers were stored in sealed Ziploc bags and kept protected from light until measurement.

### Feather colouration measurements

Yellow breast feather colouration was measured in the laboratory by XX (2013-2020) and XX (2021-2022) using a spectrophotometer (AVASPEC-2048; Avantes, Apeldoorn, The Netherlands) and a deuterium-halogen light source (AVALIGHT-DH-S lamp; range of 300-700nm; Avantes, Apeldoorn, The Netherlands). Feathers were illuminated from a 90° angle with a 200µm fibre optic probe at a distance of 2mm. To exclude ambient light, a probe mount consisting of a black rubber cap was placed on the probe. Reflectance data was computed relative to a white reference (WS-1; Ocean Optics, Dunedin, FL, USA) and a dark reference (a black felt background). Feathers were displayed on a piece of black felt providing a background reflectance of 0. Two sets of four randomly chosen feathers were superposed for each individual, reflecting the bird plumage (Quesada & Senar, 2006). We measured each set of feathers three times, with the probe removed between each measure, and used the average spectra to compute the colour variables. Average spectra obtained for males and females from both habitats are shown in Fig S2.

Based on these spectra, we computed three variables that are commonly used to describe colour variation using the R package *pavo* (v. 2.8.0; Maia et al. 2019): mean brightness, yellow chroma and UV chroma (Peneaux, Grainger, et al., 2021; Sepp et al., 2018). We calculated mean brightness as the integral area under the reflectance curve divided by the width of the interval 300-700nm, representing the overall intensity of light reflected by the feathers. Yellow chroma was estimated as the relative difference between the maximum (after the plateau at 500nm) and the minimum reflectance ([R700 – R450] / R700) and is tightly linked to the amount of carotenoids deposited on the feather (Isaksson et al., 2008). Finally, we obtained UV chroma as the proportion of the total reflectance under the ultraviolet range ([300-400] nm). UV chroma is linked both to the carotenoid content and to feather microstructure (Shawkey et al., 2006). These three variables are moderately correlated (correlation ranges from −0.40 to +0.36; see Table S1). Our measurements were strongly repeatable (ranges of repeatability estimates for: yellow brightness: [0.74 – 0.92]; yellow chroma: [0.57 – 0.9]; yellow ultraviolet chroma: [0.35 – 0.85]; see Table S2 for details).

### Climatic variables

We computed the mean temperature and rainfall for each moulting period (June 1st-September 30th), for every year between 2012 and 2021, to represent the climatic conditions experienced by great tits while renewing their plumage (Jenni & Winkler, 2020; López-Idiáquez et al., 2022). We calculated these values from daily mean temperature and rainfall data, where daily mean temperature corresponds to the average of the minimum and maximum daily temperatures. For the city, we used information from a meteorological station at the Centre d’Ecologie Fonctionnelle et Evolutive (CEFE) in Montpellier (Figure S1). For the forest, we used temperature data from the weather station in Saint Martin de Londres located at about 15 km from the forest and 25 km from the city (Figure S1). The information from this station has been successfully used in previous studies and provides a reliable measure of the temperature experienced in the forest (see Bonamour et al., 2019, S3 for further detail). For rainfall, we used data from the Montarnaud weather station, around 3 km from the forest and 15 km from the city. Both forest stations are part of the French national meteorological service (MeteoFrance, https://donneespubliques.meteofrance.fr/).

### Statistical analysis

All data analyses were performed with the software R (version 4.2.2, R Core Team, 2022). We modelled variation in the three proxies of breast colourations (brightness, yellow chroma and UV chroma) using Linear Mixed Models (LMM) with the package *lmerTest* (v. 1.1-33; Kuznetsova, Brockhoff, and Christensen 2017). LMMs described below included the following random effects: individual identity since some birds were captured several times along the study period (range [1-6], average = 1.37), year (as a categorical variable) and sites of capture (n=9, one forest site and eight urban sites that capture different degrees of urbanisation in the city sampled, see Caizergues et al., 2024) to control for the non-independence of data within years, and sites. We used the package *ggeffects* (v.1.2.3; Lüdecke 2018) to estimate marginal means and the confidence intervals.

### Urban vs rural populations comparison

We analysed the differences in colouration between urban and forest populations for each of the three colour components. Due to low correlation among the three variables (Table S1), we tested them independently by fitting three sets of LMMs with a normal distribution of errors. Full models included as explanatory variables: habitat (urban vs forest), sex (male vs female), and age (yearling vs adult). Additionally, we examined two-way interactions between habitat and sex, as well as habitat and age, based on our hypothesis that habitat effects may vary according to sex and age. We made post-hoc tests using *emmeans* package (v. 1.8.7, Lenth, 2023) to obtain contrast evaluation within interactions.

### Temporal trends in colouration

We tested for temporal trends on the three great tit breast patch components of colouration by fitting three LMMs with a normal distribution of errors, one for each colour variable. Years were scaled to a mean of zero and a standard deviation of one to account for differences in scales between colouration and years. As explanatory variables, we included year (as a continuous variable) in addition to its quadratic version (as orthogonal polynomial), to capture a concave pattern observed in graphical exploration. Habitat (urban vs forest), sex (male vs female) and age (yearling vs adult) were also included as fixed effects, along with the two-way interactions of linear and quadratic years with population, sex and age, to explore the temporal variation of the difference of colouration between populations.

### Temporal trends in climate

We analysed the temporal trends in mean temperature and rainfall during moulting period between 2012 and 2021 for each habitat using linear models with a normal distribution of errors. The dependent variables were temperature and rainfall. Year (as a continuous variable), habitat and their two-way interaction were included as explanatory variables.

### Climate association with plumage colouration

We used climatic variable (temperature and precipitation) from the year before we have the colouration samples since the moult occurs in the summer preceding the reproduction events we follow, for example the colouration of a bird captured in the reproductive season of 2014 was linked to the climatic data from the summer of 2013.

First, to explore the association between great tit colouration and climate, we fitted LMMs on data from the two habitats (urban / forest). Temperature ranges did not strongly overlap between the city and the forest population (22,3 – 23,9°C for city and 20,9 – 22,4°C for the forest) hence the climatic variables were mean centred within each habitat, to obtain relative temperatures and rainfall to compare them. We analysed the effect of climatic variables (temperature or rainfall), habitat, sex, age and the two-way interactions between climatic variables and the three other variables. This model aims at testing for a differential effect of climate between urban and forest habitats.

We then analysed each habitat separately to investigate associations between colour and climate (using climate absolute values). The climatic variables were tested in separate models. Full models included average temperature or rainfall, sex and age as explanatory variables, along with the two-way interactions between climatic variables and age and climatic variables and sex.

In all the models, we assessed the significance of fixed effects and their interaction using the Anova() function from the *car* package (Fox & Weisberg, 2019), which computes tests based on Type III sums of squares. P-values were obtained using F-tests. We reduced our models using a stepwise backwards method. We present in the results the ANOVA of full models with the steps of removal.

## Results

### Urban effect on great tit colouration in relation to sex and age

### Yellow chroma

Birds in urban environments exhibit a significant reduction in yellow chroma compared to forest birds, although this effect differs between each sex and age class (interaction habitat and sex: 0.04 ± 0.02, F_1, 1047.81_=6.5, p=0.01; interaction habitat and age: −0.07 ± 0.02, F_1, 1528.20_=20.9, p<0.001, Table S3, Fig.1A and 1B). Specifically, males in urban areas show a significantly 14% lower yellow chroma (−0.08±0.03, p=0.03) than their forest counterparts, while females do not show reduced chroma in the city (−0.04±0.03, p=0.18) compared to forest ones. In addition, sex dichromatism between males and females was only present in the city, where males show 6% less yellow chroma than females (in the city: −0.03±0.008, p<0.001, forest p = 0.62, Fig. 1A).

**Figure 1.**
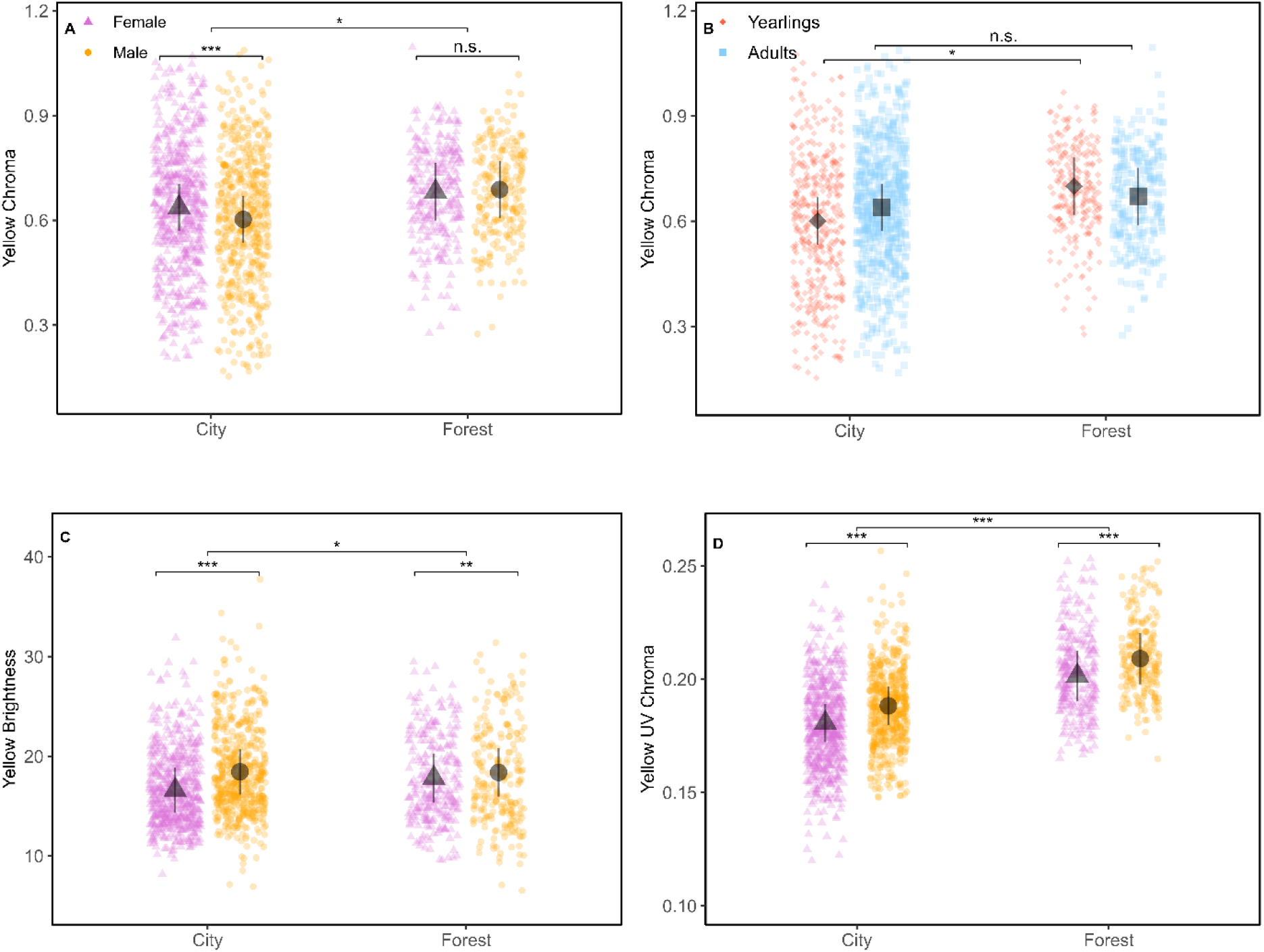
Differences in yellow breast feather colouration between habitats, and across sex and age: yellow chroma (A and B), brightness (C), UV chroma (D). Estimates and confidence intervals are outputs from the models presented in Table S3. Asterisks denote significant differences (*** p<0.001, ** p<0.01, * p<0.05) and n.s. no significant differences. In A, C and D, violet triangles represent females and orange dots males. In B red diamonds represent juveniles and blue squares represent adults.

Regarding age, the difference between the city and the forest is marginally significant for yearlings, with urban ones showing a 17% reduction in chroma compared to their forest counterparts (−0.09±0.03, p=0.02, Fig. 1B), but this effect is not observed in adults (p=0.34, Fig. 1B). Last, the age-related differences in yellow chroma are reversed between habitats: in the city, adults are 6% more chromatic than yearlings (age effect in the city: 0.04±0.009, p<0.001, Fig. 1B), whereas in the forest, yearlings show 4% higher chroma than adults (age effect in the forest: −0.03±0.01, p=0.02, Fig. 1B).

### Yellow brightness

Urban and forest birds exhibit significant differences in yellow brightness, but these patterns vary by sex (Table S3, Fig. 1C). On one hand, forest females display 7% brighter plumage than their urban counterparts (1.21±0.49, p=0.06, Fig. 1C), although this association is marginally significant. Males, on the other hand, do not show such difference (p=0.93, Fig. 1C). Sexual dichromatism is, once again, more pronounced in urban habitats, with males displaying 11% brighter yellow feathers than females (males: 1.84±0.17, p<0.001, Fig. 1C), while the difference in the forest is more subdued, with males 3% brighter than females (0.58±0.25, p=0.02, Fig.1C)

### UV chroma

We find that urban great tits show 11% lower UV Chroma compared to forest birds (forest: 0.02 ± 0.005, F_1, 5.15_=21.1, p=0.005, Table S3, Fig. 1D), with similar patterns across sex and age classes (*i.e.,* no significant interactions, Table S3).

### Temporal trends in colouration

#### Yellow chroma

We find a non-linear temporal trend in yellow chroma, with both habitats showing a minimum value in 2017 (Table 1, Fig. 2A). In addition, we find significant interactions between year^2^ and sex, as well as between year and habitat (Table 1, Fig. 2A).

**Figure 2:**
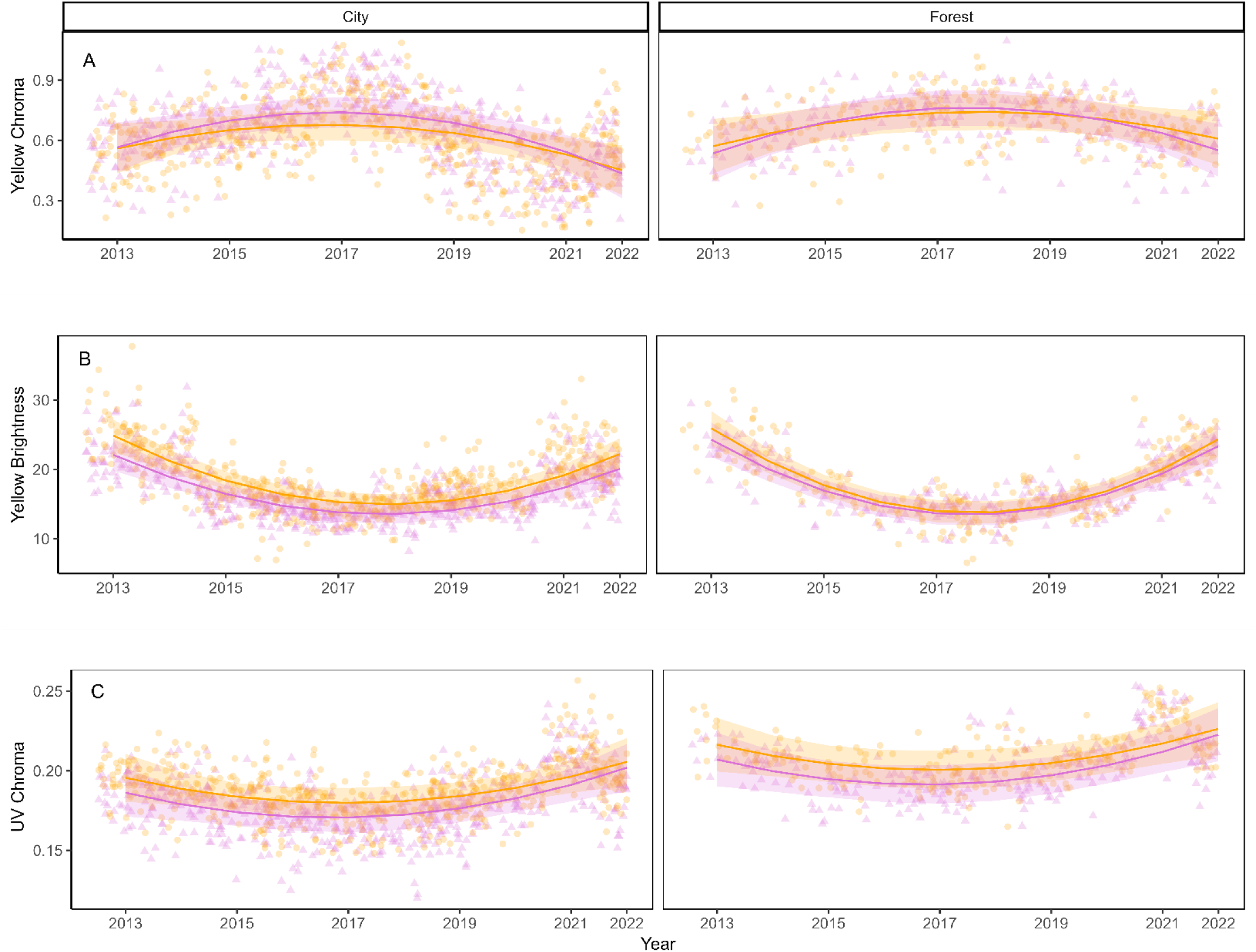
Quadratic patterns of male and female great tits yellow breast colouration in the city of Montpellier (left panels) and the forest of La Rouvière (right panels) across time (2013-2022); (A) Yellow chroma, (B) Brightness and (C) UV Chroma. The lines show the predicted slope values (Table S4). The dots represent the raw data. 95% confidence intervals are shown as the coloured stripes. Violet triangle lines represent the females, orange dots and lines the males.

**Table 1:**
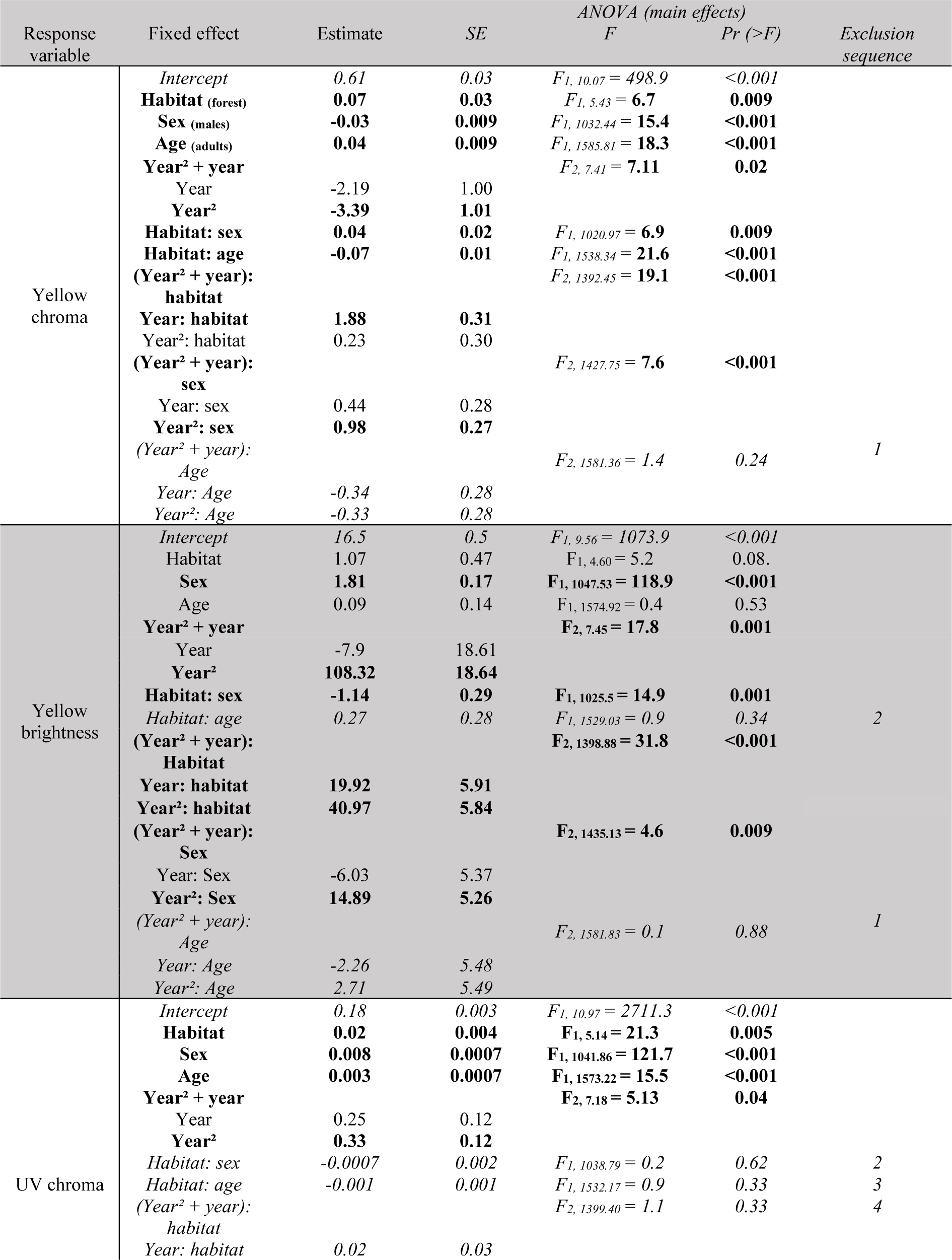
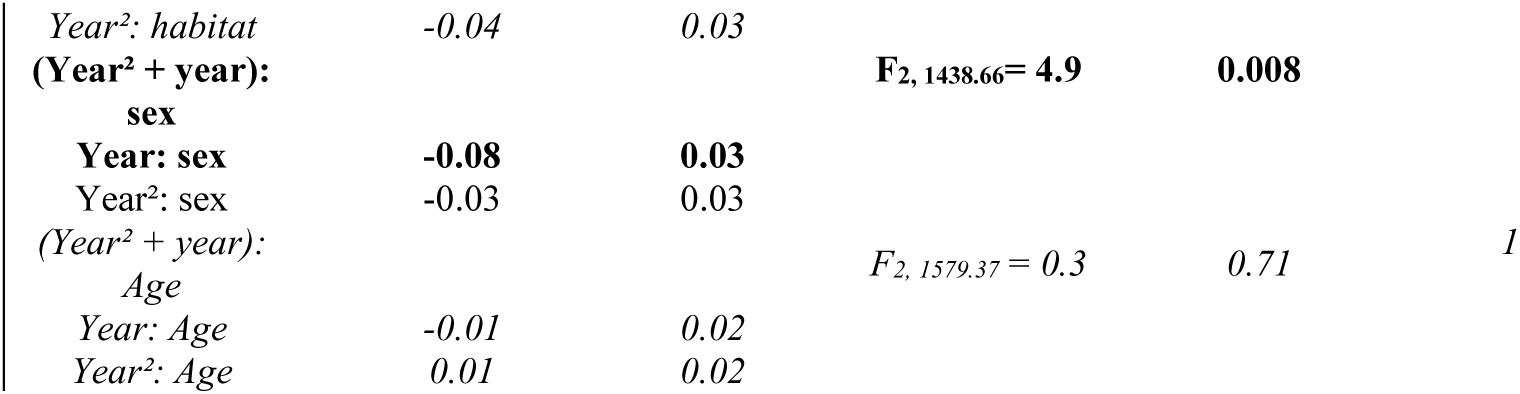
Linear and quadratic temporal trends for the colouration of urban and rural great tits. Type III sums of squares Anova are used and P-values are obtained with F-tests. Significant variables are in bold. Reference levels of habitat, sex and age are the city, females and yearlings. Number of observations for each habitat: N*_city_* = 1102 / N*_forest_* = 504. Number of individuals for each habitat: N_city_ = 811 / N_forest_ = 356. Values for excluded variables refer to the step before exclusion and are in italic. Year² + year represents the polynomial term for the linear and quadratic associations.

#### Yellow brightness

Brightness also follows a quadratic term over time, with a minimum in 2018 in both habitats (Table 1, Fig. 2B). There is a significant interaction between year² and habitat, and year² and sex (Table 1, Fig.2A).

#### UV chroma

A quadratic association is observed for UV Chroma and year, with a minimum UV colouration in 2017 (Table 1, Fig. 2C). Additionally, there is a significant interaction between the linear temporal effect and sex (Table 1, Fig. 2C).

### Temporal trends in climate

Our analyses reveal that urban areas are on average 1.4°C warmer than the forested area nearby (city as reference: −1.4±0.21, F_1, 17_=41.9, p<0.001, Fig.3A). There is also a significant increase in temperature over time, with a rise of 0.08°C per year (0.08±0.04, F_1, 17_ =5.1, p=0.03, Fig. 3A). No significant difference in rainfall is observed between the two habitats (0.46±0.49, F_1, 17_=0.87, p=0.37, Fig.3B), nor is there a significant temporal trend in rainfall across the study decade (−0.07±0.09, F_1, 17_=0.69, p=0.42, Fig.3B).

**Figure 3:**
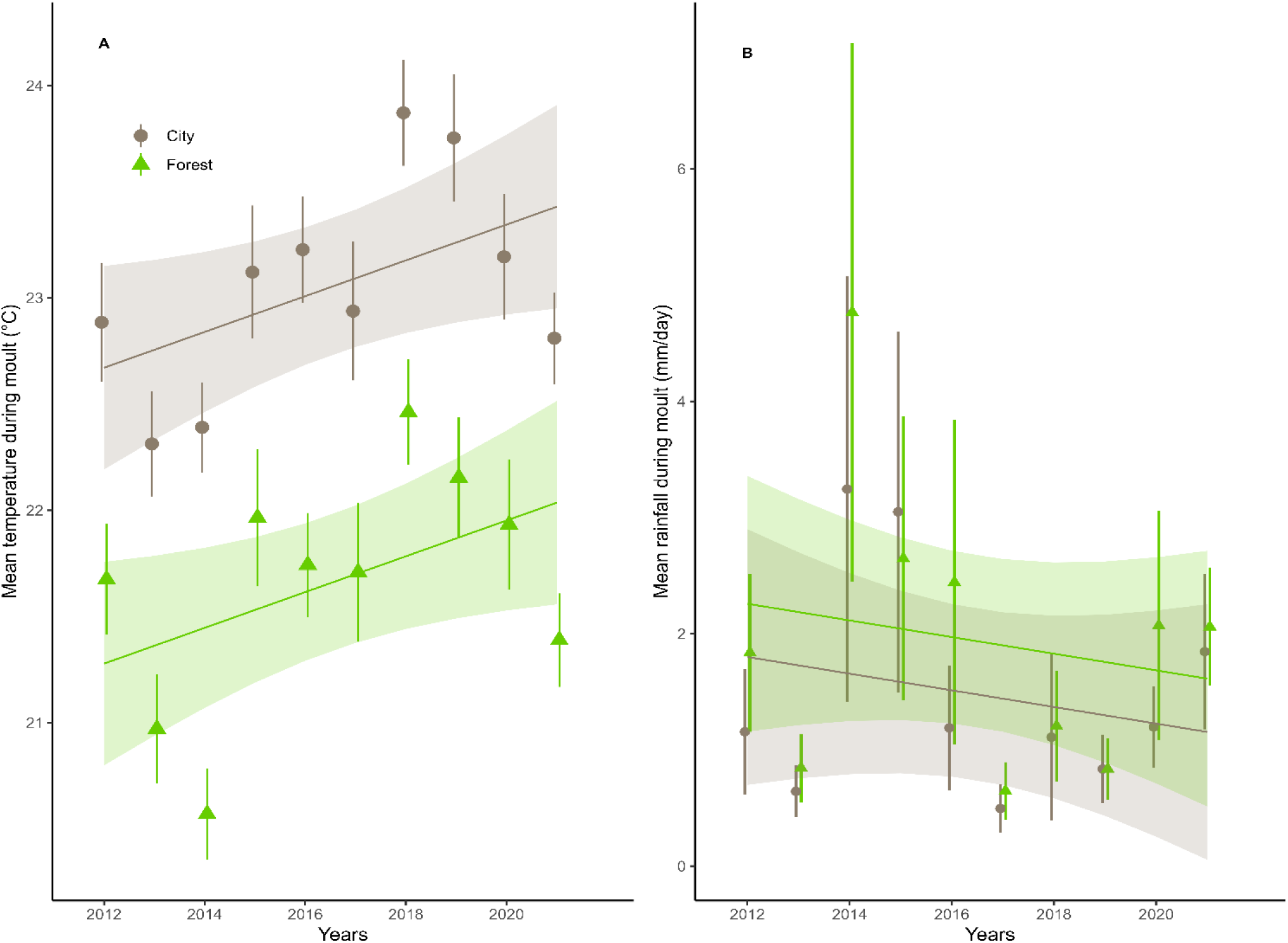
Temporal trends in mean temperature (A) and mean rainfall (B) during great tit moulting over a decade (2012-2021). The lines show the slope predicted by models presented in ‘*Temporal trends in climate’*. 95% confidence intervals are shown as the coloured stripes. The dots show the mean temperature and rainfall for the moulting period each year and whiskers represent the standard errors. Green triangles represent observations from the forest and grey dots from the city.

### Climate association with colour

When modelling the effects of relative temperatures and rainfall on colouration, we find that yellow chroma is more negatively impacted by higher relative temperature in the city (−0.19 or 30% chroma for an augmentation of 1°C) than in the forest (−0.08 or 10 % chroma for an augmentation of 1°C) (Table 2; Fig. 4). In contrast, yellow brightness tends to be more negatively impacted by relative temperature at moult in forest birds, where it declines more sharply with increases in temperatures (−2.74 brightness or 15% for an augmentation of 1°C) than in the city (−1.66 brightness or 10 % for an augmentation of 1°C, Table 2; Fig. 4). Furthermore, relative rainfall at moult interacts with habitat to affect both brightness and UV chroma brightness being more positively influenced by rainfall in the forest (+0.45 or 2.5% brightness for an augmentation of 1mm) than in the city. (+0.02 brightness or 0.3% for an augmentation of 1mm, Table 2, Fig. 4) while it is the contrary for UV chroma (+0.005 UV or 2.5% in the city and +0.0005 or + 0.1% in the forest for an augmentation of 1mm, Table 2, Fig.4)

**Figure 4:**
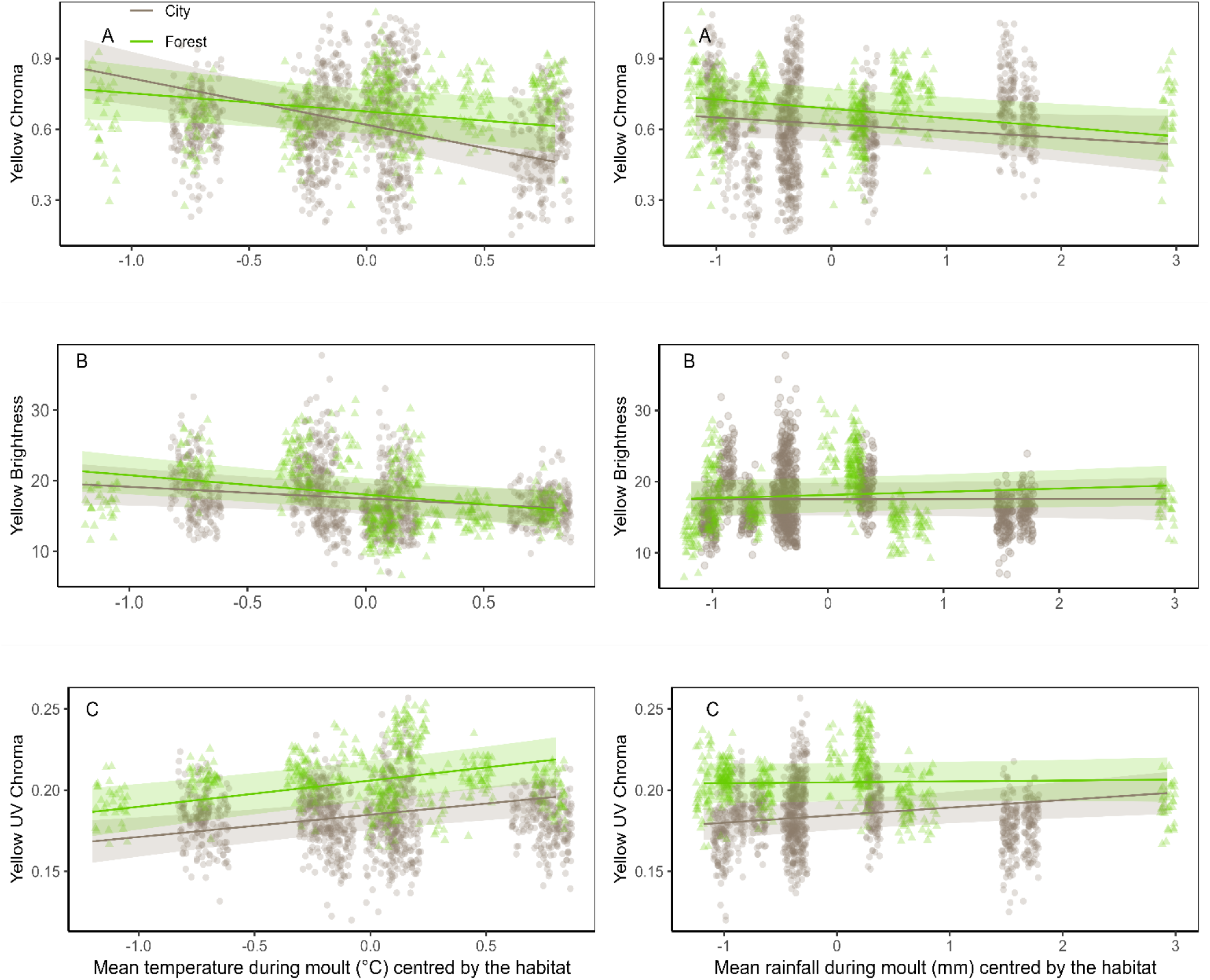
Association for great tits yellow breast colouration of the forest and the city with relative temperature and relative rainfall during the moulting period, i.e., mean temperatures (top) and rainfall (bottom) both centred by habitat; (A) Yellow chroma, (B) Yellow brightness and (C) UV chroma. Lines show the predicted slope values (Table S4). The dots show the raw data. Green triangles and lines represent the forest, grey points and lines are for the city.

**Table 2:**
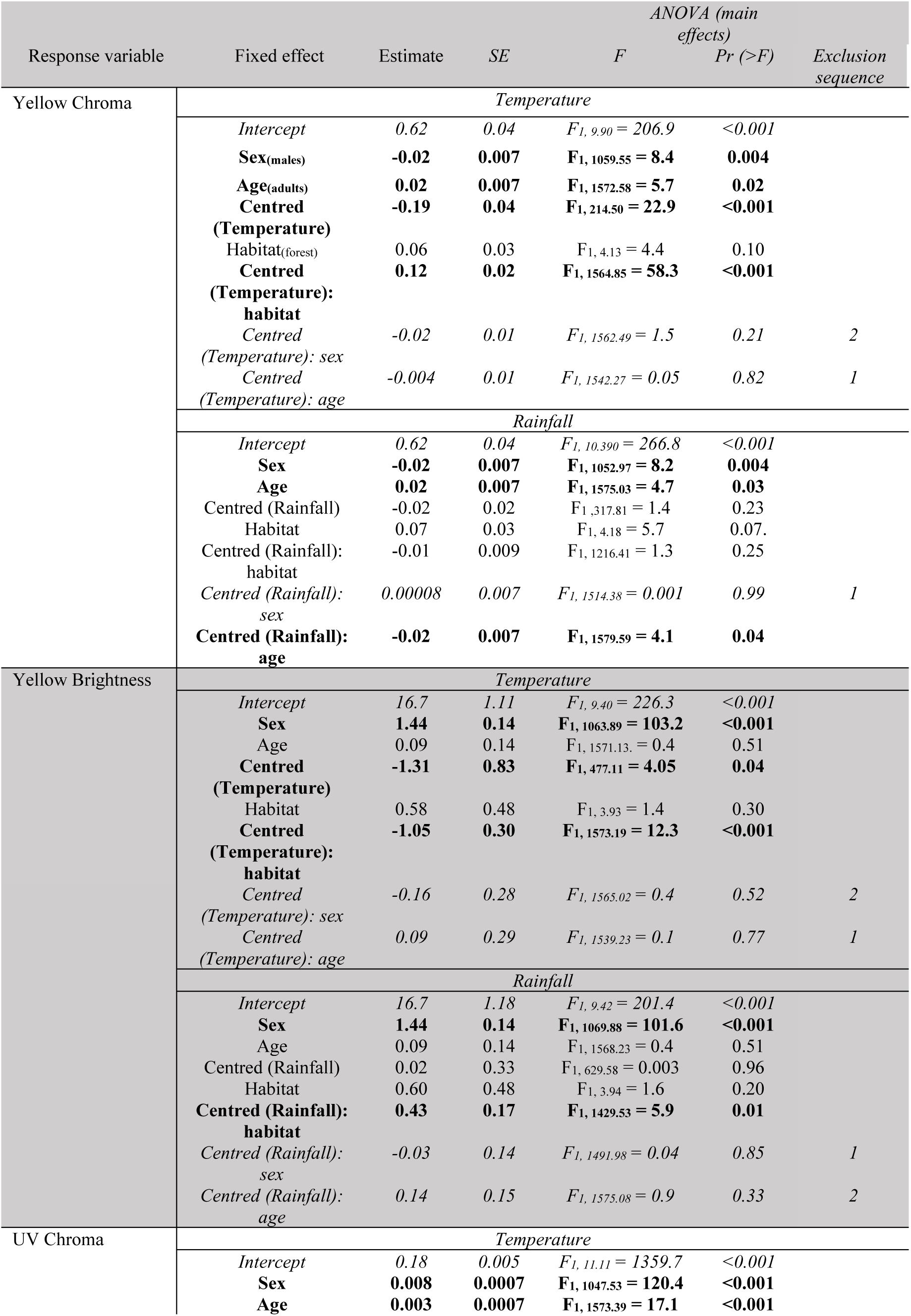
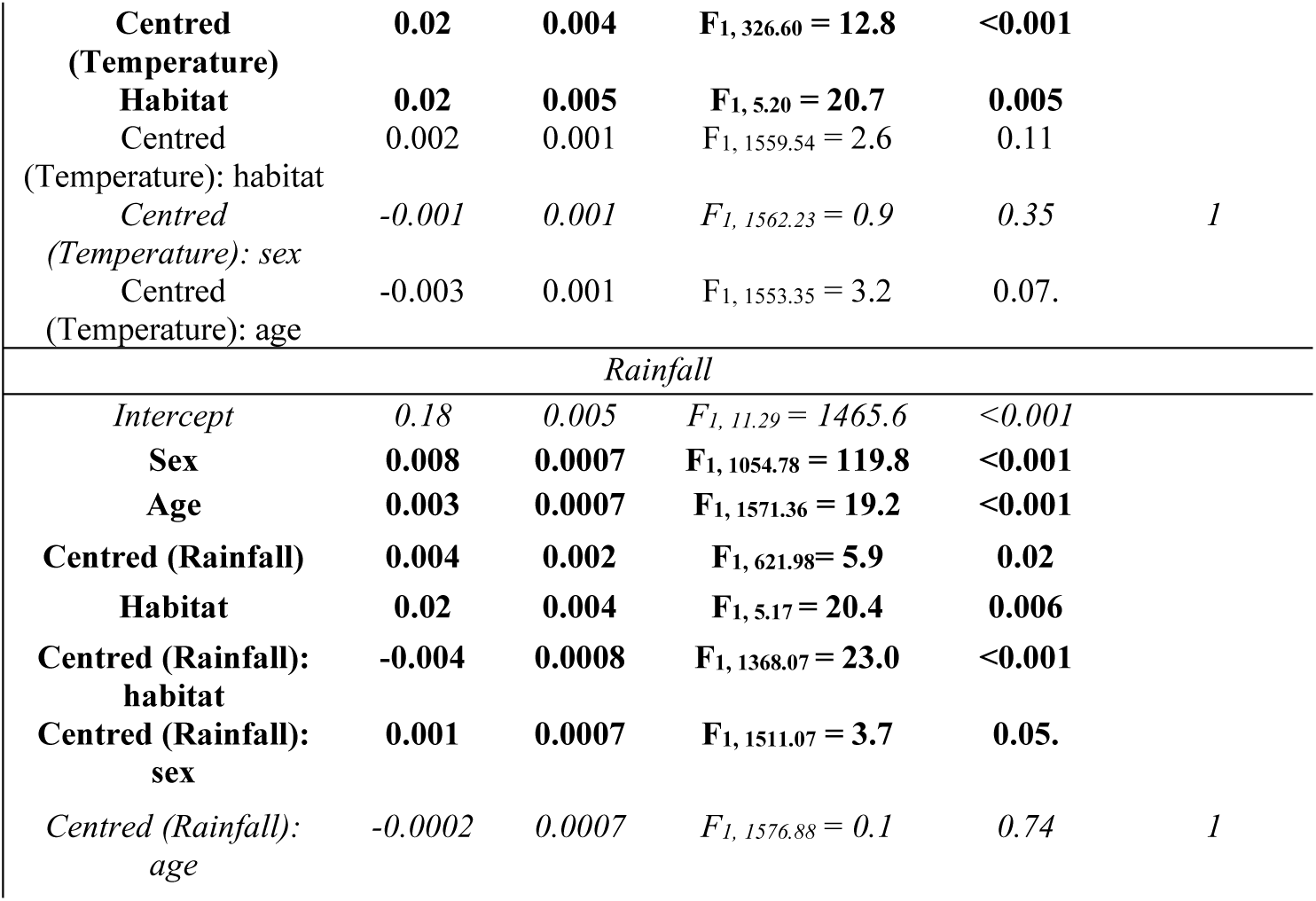
Association between climatic variables and each colouration trait of city and forest great tits, with an interaction between habitat of origin and mean temperature or rainfall, centred by the habitat. Type III sums of squares Anova are used and P-values are obtained with F-tests. Significant fixed effects are in bold. Reference levels of habitat, sex and age are city, females and yearlings. Number of observations for each habitat: N*_city_*=1102 / N*_forest_*=504. Number of individuals for each habitat: N_city_ = 811 / N_forest_ = 356. Values for excluded variables refer to the step before exclusion and are in italic.

When analysing the effects of absolute temperature and rainfall on colouration measures within each habitat, we find no significant direct effect of average temperature or average rainfall during moult on great tit colouration (Fig. S3, Table S4, range of p-values [0.22 – 0.96]). However, significant interactions between climatic variables and sex or age are observed, especially in urban habitat. For instance, urban males show a greater decline in yellow chroma with increasing temperature compared to females (F_1, 1055.24_ =4.1, p=0.04, Table S4, Fig. S3), while yearlings are more negatively affected by temperature than adults (F_1, 1084.38_ =4.7, p=0.03, Table S4, Fig. S3). For the other interactions details, see Tables S4, S5 and Figures S3, S4.

## Discussion

Our study shows that urban great tits display paler and duller colourations than their forest counterparts (around 10% less chromatic males and 7% less bright females). This pattern aligns with existing evidence supporting the “urban dullness” phenomenon and extends it by demonstrating that cities affect not only the chromatic but also the structural component of the colour, (i.e., brightness and ultraviolet components). In addition, we uncover sex- and age-dependent associations, with urban birds displaying higher sexual dichromatism compared to their forest counterparts, and yearlings displaying duller colourations than adults only in urban environment. By measuring colour and climate over a decade in these populations, we observe temporal changes in colouration alongside an increase in temperature. Furthermore, the strength of the associations between climate and colour varies between the forest and the city, being stronger in the latter. This climate-colour association also differs between sex and age classes for some colour variables. These findings highlight differential climatic impacts on colouration traits across habitats, with urban birds - especially males - being more sensitive to urban heat. We discuss the potential consequences of these changes below.

### City affects all components of colouration

The increased dullness - *i.e.,* the decrease in the chromatic component of the colouration, yellow chroma - of urban great tits compared to their forest counterparts aligns with previous studies, including a meta-analysis on several pairs of urban and rural populations across Europe (Salmón et al., 2023). It also mirrors changes observed in other avian species, such as northern cardinals (*Cardinalis cardinalis)* (Jones et al., 2010, but see Baldassarre et al., 2022 for contrasting results) and house finches (*Haemorhous mexicanus)* (Sykes et al., 2021), showing that urban bird dullness is a widespread phenomenon among species that use colour for communication (Janas et al., 2024; Leveau, 2021). Colours serve multiple signalling functions, such as species and individual recognition, as well as individual quality assessment in competitive and mating context (Hill & McGraw, 2006). How birds communicate with reduced colouration is unknown, especially since cities also affect their other main communication channel: song (Brumm et al., 2021; Halfwerk et al., 2011; Slabbekoorn & Den Boer-Visser, 2006). Given the importance of communication for social interactions – and the prediction that by 2050, two-thirds of the human population will live in cities, and exert strong impacts on the natural environment (Seto et al., 2012) – exploring how animals communicate in anthropised environments, and the consequences this may have for their reproduction and survival, is of the highest relevance.

The reduced yellow chroma in urban environments can be explained by two main environmental factors common to species that rely on carotenoids in their signalling. A diet lacking carotenoids, either due to quality or quantity, may account for the increased dullness observed in urban great tits. For example, carotenoid-rich insects are less available during reproduction in cities than in forests (Jensen et al., 2022; Pollock et al., 2017; Sinkovics et al., 2023), and fruits consumed during moult (Herrera, 1984) likely differ across habitats due to variations in plant species composition. Alternatively, a non-exclusive explanation is that urban stressors (e.g., increased pollution) reduce carotenoid availability for colour production. Carotenoids play an important role as immune-enhancing antioxidants (Pérez-Rodríguez et al., 2010); therefore, urban environments and their concomitant stressors may impose a stronger greater pressure on birds to allocate carotenoids to maintain homeostasis, rather than colouration.

Our study reveals that a new dimension of the colour can be changed in the city: the achromatic part of the colouration is reduced by approximately 10 % (brightness and UV), raising intriguing questions about the origin of this result. Urban great tits are less bright and have reduced UV chroma compared to their forest counterparts. This finding parallels observations that great tits breeding close to roads show reduced UV chroma (Grunst et al., 2020, but see Biard et al., 2017; Salmón et al., 2023 that found no change in brightness in urban areas). Brightness and UV chroma reflect the internal keratin structure and organisation of feathers, including the density and orientation of barbs and barbules (Shawkey & Hill, 2005). Reduced structural components of the feathers can be linked to factors such as habitat transformation and pollution, which can lead to a lower-quality diet and limited resources available for developing high-quality feathers. Pollution can also have a more direct effect on structural components. Particle deposition on feathers can reduce feather reflectance because of their light-scattering and absorbing properties (*e.g.,* overall reflectance and UV, Ellis et al., 2023; Griggio et al., 2011). Additionally, urban areas can impact moult duration and timing, with urban birds possibly moulting earlier (Hutton et al., 2021), resulting in fewer resources available for moult, especially if it overlaps with breeding (Svensson & Nilsen, 1997).

The decrease in chromatic yellow coloration we find is perceptible to us (see Fig.S5) and is likely noticeable to birds as well, who can detect smaller colour differences than humans. For instance, gradient changes in carotenoid-based colouration are perceptible to Zebra finches (*Taeniopygia guttata*) (Caves et al., 2018), and many birds can detect dichromatism that is invisible to us (Eaton, 2005). Further research is needed to explore how these colour changes affect bird interactions. However, it is already evident from this and other studies that carotenoid-based colouration could serve as a bioindicator (Lifshitz & St Clair, 2016; Peneaux, Hansbro & Griffin, 2021) and signal a shift in life history strategies in urban animals, prioritising traits that favour reproduction and survival over those linked to mating success (Hutton & McGraw, 2016).

### City affects differently birds depending on their age and sex

Age and sex are known to be associated with different values of coloration and to be impacted differently by environmental factors (Salmón et al., 2023; Sykes et al., 2021). In line with this hypothesis of varying sensitivity, we find different results across age and classes.

#### Age dimorphism

We find that yearlings, but not adults, exhibited a 17% decrease in chromatic colouration in the city compared to the forest. This could be due to greater sensitivity in yearlings and/or higher selective disappearance. From a mechanistic perspective, yearlings may be more sensitive as they are more likely to be outcompeted for high-quality or larger food resources (Heise & Moore, 2003), and experience faster moults, which is associated with less chromatic yellow (Ferns & Hinsley, 2008) compared to adults. While these factors apply to both urban and forest birds, their effects may be enhanced in cities, where carotenoids and other resources are scarcer (Seress et al., 2018). The pattern we find could also reflect the conditions yearlings faced as nestlings, since their moult occurs after fledging (Bojarinova et al., 1999), and urbanisation can impact nestling conditions that affect moulting conditions. Alternatively, a non-exclusive explanation is that selection against paler yearlings may be stronger in the city than in the forest. Since colour is a quality signal, yearling quality could be lower in urban areas because of the scarcity of resources and the effects of urbanisation on nestlings. This idea aligns with previous results from the study area showing that urban yearlings have lower survival rates than their forest counterparts (Caizergues et al. 2022). Analyses of Capture-Mark-Recapture data, considering colour as a covariate, will allow testing of this latter possibility.

#### Sexual dimorphism

Our results also show a change in sexual dichromatism between urban and forest birds, with urban males being 6% less chromatic and 11% brighter than females. This pattern is not observed in forest, where males and females present similar colouration. Two non-exclusive explanations can be proposed. (i) The differences between males and females can be due to higher sensitivity in urban males, leading to a greater decrease in colouration. Males may lose more colour in harsher urban conditions or experience higher mortality when investing too heavily in sexual secondary traits. (ii) The increased chroma of urban females relative to males could result from stronger sexual selection pressures on females in cities, either through female-female competition or mate choice. Theoretical models, supported by a comparative analysis (Fargevieille et al., 2023), suggest that females colouration tends to increase relative to males when parental care increases, particularly when females invest less in reproduction. Both conditions are met in urban areas, as females lay smaller clutches (Chamberlain et al., 2009; Sepp et al., 2018) and parents contribute more to feeding nestlings (Isaksson & Andersson, 2007; Jarrett et al., 2020). Nonetheless, the specific mechanisms underlying this potential increase in sexual selection on urban females remain unclear. Studies on blue tits (*Cyanistes caeruleus*) and other species emphasise the importance of female ornamental colourations as quality signals, supporting their role as secondary sexual traits (Doutrelant et al., 2020), which warrants further exploration in urban environments. Previous research on sexual dichromatism in urban environments has not reported changes in sexual dichromatism in urban compared to non-urban (Dauwe & Eens, 2008; Isaksson et al., 2005), but, overall, too few studies explicitly investigate both sexes or females (Janas et al., 2024). Further research into the causes and consequences of these shifts in sexual signals is critical. Given the link between sexual selection and adaptive potential (Gómez-Llano et al., 2021; Whitlock & Agrawal, 2009), it is crucial to assess how sexual selection operates in urban areas and whether sexual signals are still used in the same way, as environment may alter the benefits and costs of mate choice (Cronin et al., 2022).

### Biological and evolutionary implications of divergence in colouration

The divergence in colouration between urban and forest great tits, approximately a 10% difference, could carry biological and evolutionary implications. Such changes are expected to influence natural and sexual selection processes, given the role of colouration in mate choice and competition for resources (Svensson & Wong, 2011; Weaver et al., 2018). For instance, differences in dichromatism suggest that sexual selection could be relaxed in males or heightened in females in urban environments, potentially affecting mate selection and reproductive success. However, it is to keep in mind that signalling values of colouration are relative to the local phenotype and environment. For example, a study showed that female preference is plastic and can adjust to the available phenotype in urban areas (Giraudeau et al., 2018). In addition, a recent study in urban great tits showed that paler urban individuals tended to have higher reproductive success than yellower urban individuals, indicating that duller urban colours could be reflective of an adaptive responses to the urban environment (Bekka et al., 2025). Additionally, sexual imprinting could reinforce these patterns, as urban birds reared by parents and surrounded by individuals with duller phenotypes may develop a preference for birds displaying the local phenotype. Supporting this hypothesis, a study revealed genetic differentiation between urban and forest. populations in Montpellier (Perrier et al., 2018) that could be a cause and/or a consequence of colour divergence. Moreover, this shift extend to other traits, such as the black-tie colouration of great tits (Dauwe & Eens, 2008; Senar et al., 2014, Sandmeyer et al. *under review*) and songs (Halfwerk et al., 2011; Slabbekoorn & Den Boer-Visser, 2006), which also change in urban areas. This suggests that multiple secondary sexual traits are shaped by urban pressures, reshaping urban bird phenotypes and potentially affecting gene flow and local adaption. In addition, the chromatic contrast between the bird’s plumage and its background likely differs between urban and forest habitats. More achromatic and grey urban environments may enhance the detectability of bright plumages, while green and heterogeneous forest backgrounds may favour more saturated colours. Thus, duller plumage in urban birds could also be the result of adaptive background matching to reduce predation risk or modify visual signalling efficiency (Fialko & Price, 2025; Gomez & Théry, 2007). Testing this hypothesis using visual modelling is an interesting question for future research. The consequences of these shifts in selection pressures could alter population dynamics and adaptation strategies, ultimately impacting the evolutionary trajectory of these birds (Charmantier et al., 2024). Our study provides important insights into how anthropogenic factors reshape avian colouration and calls for a deeper understanding of the causes and implications for ecological and evolutionary processes.

### Temporal trends in colouration

We observe a similar quadratic pattern in temporal trends in both habitats, with mean colouration being higher (chroma) or lower (brightness and UV chroma) between 2016 and 2018 than in the years before and after, delineating three distinct periods: before 2016, between 2016 and 2018, and after 2018. Such temporal variation could arise from environmental (*e.g.,* climate change) and/or demographic processes. To investigate this unexpected pattern, we examined potential demographic drivers such as age structure (*e.g.,* variations in the proportion of yearlings) and population turnover due to immigrant arrivals. However, we found no consistent year-to-year variation in either habitat (Fig. S6, Table S6), therefore these factors do not explain the observed trends. One possible alternative explanation could be regional resource pulses affecting both forest and urban areas. Further investigation is needed to identify the environmental factors driving these temporal changes in colouration. Previous research on temporal trends in colouration reported linear changes in colouration linked to environmental factors such as temperature (Evans & Gustafsson, 2017; López-Idiáquez et al., 2022). We also explored the role of temperature as a driver of the temporal trends, but find no associations to explain the quadratic pattern. Furthermore, our analysis shows that the dullness of urban birds compared to forest birds remains consistent over time. Previous studies focus on colouration differences have typically focused on one- or two-year period, which limits their ability to investigate the consistency of such patterns over longer timeframes. Since our findings indicate that the colouration divergence between habitats has neither emerged in the last decade nor is changing, we propose that is it likely a result of long-term urbanisation effects rather than recent changes, and that colouration difference remains relatively stable over time.

### Deciphering urbanisation and climate effects on colouration

Interactions between climate and the effects of urbanisation have been rarely investigated. Our results suggest differences in the climate sensitivity of great tit colouration between urban and forest habitats. Overall, an increase in temperature and precipitation has a stronger negative effect on the chromatic colour variable in the city than in the forest (−30% for +1°C in the city versus −10% in the forest), while they have a stronger positive effect on structural variables in the forest than in the city (2.5% of brightness and UV in the forest for +1mm of rain versus 0.1 and 0.3% respectively, in the city). This urban-specific climate effect could reflect environmental conditions prevailing in cities (e.g., higher temperature due to urban heat island effect), and thus a greater sensibility of urban birds. This result also suggests that carotenoid availability and assimilation by great tits could be influenced by urban conditions combined with higher temperatures. An interesting parallel can be made with a previous study in blue tits, where stronger climate-colour associations were reported in hotter, drier Corsica compared to the mainland (López-Idiáquez et al., 2022). Such a result can also be expected for the colour of ectotherms, such as insect, fish and reptiles. For example, a previous study found similar effects on butterflies (Diamond et al., 2014). While the observed correlation between climate and colour in this study arises from the interaction of habitat and climate, our results suggest that urbanisation could amplify climate change effects, imposing additional pressures on urban-dwelling individuals. Although our findings are based on a single urban-forest pair, this study highlights a promising avenue for future research, and further studies should assess the extent to which the pattern observed applies across multiple populations, species, and climatic gradients.

## Conclusion

In conclusion, our study highlights the impact of urbanisation on both the structural and chromatic components of carotenoid-based colouration, and shows that ages and sexes are not equally affected. Furthermore, we provide evidence of interactive effects between climate and habitat on carotenoid-based colouration, impacting differently urban and rural birds. However, the observed effects of climate on colouration call for larger than 10-year temporal scale confirmation, particularly in investigating its interaction with the urban heat island phenomenon. Additionally, our findings underscore a significant gap in the literature regarding the experience of urban-dwelling birds during the moulting period, including the timing of moult, die, and health during this critical phase of the year. Further research is also needed to investigate how changes in ornamental colouration affect animal social interactions and sexual selection dynamics in urban environments. Specifically, it is important to determine whether modified signals in urban contexts can still convey the same information to conspecifics and reliably indicate the quality of the signal bearer. Given the rapid expansion of urban areas in the coming years and the escalating impact of climate, addressing these questions is both timely and essential.

## Authors contribution

Anne Charmantier, Claire Doutrelant and Arnaud Grégoire conceived and funded the study. Anne Charmantier, Claire Doutrelant, Amélie Fargevieille, Pablo Giovannini, Arnaud Grégoire, and Samuel Perret collected field data. Maria del Rey and Amelie Fargevieille measured colours in the lab. David López-Idiáquez and Lisa Sandmeyer developed the statistical methodology. Lisa Sandmeyer performed the statistical analyses. Lisa Sandmeyer wrote the first draft and the manuscript. All the authors contributed to the draft and gave final approval for publication.

## Acknowledgements

We deeply thank the numerous students, technicians and researchers who have contributed to the long-term monitoring, especially Annick Lucas, Christophe de Franceschi, Marcel Lambrechts, Samuel Caro, Céline Teplitsky, Virginie Demeyrier, Aude Caizergues and Megan Thompson. Additionally, we thank the GNAUM, the Montpellier City Council, the Montarnaud City Hall and the Zoo du Lunaret for their support. We also thank Aleksandra Walczyńska, Caroline Isaksson and three anonymous reviewers for helpful comments on a previous version of the paper. Preprint version 6 of this article has been peer-reviewed and recommended by Peer Community in Ecology (https://doi.org/10.24072/pci.ecology.100800, Walczyńska, 2025).

## Fundings

This study is part of the long-term Studies in Ecology and Evolution (SEE-Life) program of the CNRS, with long-term monitoring funded by OSU-OREME. It was also supported by the ANR (France grants to A.C., Project URBANTIT ANR-19-CE34-0008-05, project ACACIA ANR-AAPG-2022-252886).

## Conflict of interest disclosure

The authors declare no financial conflict of interest.

## Data, script, and supplementary information availability

All data and scripts are available via a Zenodo repository: https://doi.org/10.5281/zenodo.12657179.

## Notes

### Competing Interest Statement

The authors have declared no competing interest.

### Summary of Updates

This version now have a link to the revision process with PCI Ecology and the recommandation.

