## supplementary information for "Disentangling urbanisation, climate effects and their interaction on ornamental colourations"

Short title: Colouration, urbanisation and climate

**Table S1**: Coefficient of correlations and 95% confidence intervals between the three colouration traits for the whole dataset (1606 colour measures for each trait). Each of these correlations is significant ($p < 0.001$) based on the Pearson test.

| | Yellow Chroma | Yellow Brightness | Yellow UV Chroma |
|---|---|---|---|
| Yellow Chroma | - | -0.41 [-0.45, -0.36] | -0.36 [-0.40, -0.32] |
| Yellow Brightness | -0.41 [-0.45, -0.36] | - | 0.36 [0.32, 0.40] |
| Yellow UV Chroma | -0.36 [-0.40, -0.32] | 0.36 [0.32, 0.40] | - |

**Table S2**: Repeatability for each colouration variable among populations and for each year, using the within individual variation across measures. Number of measures represent the 6 spectra taken from each individual (1 per spectrum).

| | Yellow Chroma | Yellow Brightness | Yellow UV Chroma | Number of measures | Number of individuals | Yellow Chroma | Yellow Brightness | Yellow UV Chroma | Number of measures | Number of individuals |
|---|---|---|---|---|---|---|---|---|---|---|
| Years | | | *Forest* | | | | | *City* | | |
| 2013 | 0,84 | 0,84 | 0,81 | 708 | 118 | 0,78 | 0,85 | 0,77 | 228 | 38 |
| 2014 | 0,82 | 0,85 | 0,78 | 678 | 113 | 0,73 | 0,85 | 0,62 | 366 | 61 |
| 2015 | 0,74 | 0,74 | 0,71 | 661 | 111 | 0,68 | 0,74 | 0,61 | 300 | 50 |
| 2016 | 0,82 | 0,9 | 0,83 | 896 | 161 | 0,7 | 0,85 | 0,78 | 273 | 49 |
| 2017 | 0,79 | 0,81 | 0,78 | 726 | 130 | 0,66 | 0,89 | 0,68 | 417 | 71 |
| 2018 | 0,86 | 0,91 | 0,85 | 780 | 133 | 0,88 | 0,91 | 0,84 | 356 | 63 |
| 2019 | 0,85 | 0,86 | 0,78 | 1084 | 188 | 0,72 | 0,75 | 0,69 | 408 | 68 |
| 2020 | 0,9 | 0,83 | 0,78 | 648 | 118 | 0,75 | 0,84 | 0,71 | 375 | 65 |
| 2021 | 0,8 | 0,9 | 0,61 | 945 | 158 | 0,57 | 0,88 | 0,35 | 471 | 79 |
| 2022 | 0,82 | 0,91 | 0,63 | 864 | 146 | 0,73 | 0,92 | 0,57 | 513 | 87 |

**Table S3:** Comparison of urban and rural great tits' yellow breast colouration. Type III sums of squares Anova are used and P-values are obtained with F-tests. Significant fixed effects are in bold. The effect of the habitat was estimated by taking the city as a reference. The effect of sex was estimated by taking the females as a reference and the effect of age by taking yearlings as a reference. Number of observations for each habitat: $N_{city}$ = 1102 / $N_{forest}$ = 504. Number of individuals for each habitat: $N_{city}$ = 811 / $N_{forest}$ = 356. Values for excluded variables refer to the step before exclusion and are in italic.

| | | | | *ANOVA (main effects)* | | |
|---|---|---|---|---|---|---|
| | Fixed effect | Estimates | *SE* | *F* | *Pr (>F)* | *Exclusion sequence* |
| Yellow chroma | *Intercept* | *0.62* | *0.03* | *$F_{1, 11.41}$ = 311.7* | *<0.001* | |
| | **Habitat (forest)** | **0.08** | **0.03** | **$F_{1, 5.40}$ = 7.1** | **0.04** | |
| | **Sex (males)** | **-0.03** | **0.009** | **$F_{1, 1060.65}$= 14.7** | **<0.001** | |
| | **Age (adults)** | **0.04** | **0.009** | **$F_{1, 1598.01}$ = 19.8** | **<0.001** | |
| | **Habitat: sex** | **0.04** | **0.02** | **$F_{1, 1047.81}$ = 6.5** | **0.01** | |
| | **Habitat: age** | **-0.07** | **0.02** | **$F_{1, 1528.20}$ = 20.9** | **<0.001** | |
| Yellow brightness | *Intercept* | *16.6* | *1.17* | *$F_{1, 9.58}$ = 199.6* | *<0.001* | |
| | Habitat | 1.21 | 0.49 | $F_{1, 4.62}$ = 5.9 | 0.06. | |
| | **Sex** | **1.84** | **0.17** | **$F_{1, 1066.08}$ = 115.8** | **<0.001** | |
| | Age | 0.11 | 0.14 | $F_{1, 1575.82}$ = 0.6 | 0.42 | |
| | **Habitat: sex** | **-1.27** | **0.30** | **$F_{1, 1040.28}$ = 17.5** | **<0.001** | |
| | *Habitat: age* | *0.28* | *0.29* | *$F_{1, 1522.61}$ = 0.90* | *0.34* | *1* |
| UV chroma | *Intercept* | *0.18* | *0.004* | *$F_{1, 12.09}$ = 1745.9* | *<0.001* | |
| | **Habitat** | **0.02** | **0.005** | **$F_{1, 5.15}$ = 21.2** | **<0.001** | |
| | **Sex** | **0.008** | **0.0007** | **$F_{1, 1056.16}$ = 119.3** | **<0.001** | |
| | **Age** | **0.003** | **0.0007** | **$F_{1, 1573.72}$ = 15.5** | **<0.001** | |
| | *Habitat: sex* | *-0.001* | *0.002* | *$F_{1, 1048.42}$ = 0.5* | *0.49* | *1* |
| | *Habitat: age* | *-0.001* | *0.001* | *$F_{1, 1534.72}$ = 0.9* | *0.33* | *2* |

**Table S4:** Table of the association between climatic variables and each colouration trait of city and forest great tits. Type III sums of squares Anova are used and P-values are obtained with F-tests. Significant fixed effects are in bold. Reference levels of sex and age are females and yearlings. Number of observations for each habitat: $N_{city}$ = 1102 / $N_{forest}$= 504. Number of individuals for each habitat: $N_{city}$ = 811 / $N_{forest}$ = 356. Values for excluded variables refer to the step before exclusion and are in italic.

| Response variable | Fixed effect | Estimates | *SE* | *ANOVA (main effects)* *F* | *Pr (>F)* | *Exclusion sequence* | Estimates | *SE* | *ANOVA (main effects)* F | *Pr (>F)* | *Exclusion sequence* |
|---|---|---|---|---|---|---|---|---|---|---|---|
| Yellow Chroma | | | | *City* | | | | | *Forest* | | |
| | *Intercept* | *1.25* | *2.08* | *$F_{1, 8.17}$ = 0.36* | *0.55* | | *-0.64* | *0.93* | *$F_{1, 8.09}$ = 0.5* | *0.49* | |
| | **Sex (males)** | **0.80** | **0.41** | **$F_{1, 1055.24}$ = 3.8** | **0.05.** | | 0.006 | 0.01 | $F_{1, 322.99}$ = 0.3 | 0.59 | |
| | **Age (adults)** | **0.03** | **0.009** | **$F_{1, 1087.03}$ = 14.2** | **<0.001** | | **-0.02** | **0.01** | **$F_{1, 466.68}$ = 4.4** | **0.04** | |
| | Temperature | -0.03 | 0.09 | $F_{1, 8.17}$ = 0.1 | 0.77 | | 0.06 | 0.04 | $F_{1, 8.09}$ = 2.0 | 0.19 | |
| | **Temperature : sex** | **-0.04** | **0.02** | **$F_{1, 1055.24}$ = 4.1** | **0.04** | | *0.02* | *0.02* | *$F_{1, 483.83}$ = 0.86* | *0.35* | *2* |
| | *Temperature: age* | *0.002* | *0.02* | *$F_{1, 1069.42}$ = 0.02* | *0.90* | *1* | *-0.01* | *0.02* | *$F_{1, 473.15}$ = 0.3* | *0.60* | *1* |
| | *Intercept* | *0.57* | *0.08* | *$F_{1, 8.50}$ = 46.7* | *<0.001* | | *0.70* | *0.05* | *$F_{1, 8.27}$ = 197.8* | *<0.001* | |
| | **Sex** | **-0.03** | **0.009** | **$F_{1, 701.12}$ = 14.1** | **<0.001** | | 0.006 | 0.01 | $F_{1, 323.07}$ = 0.3 | 0.59 | |
| | **Age** | **0.07** | **0.02** | **$F_{1, 1086.47}$ = 14.4** | **<0.001** | | **-0.02** | **0.01** | **$F_{1, 465.27}$ = 4.5** | **0.03** | |
| | Rainfall | 0.04 | 0.05 | $F_{1, 8.29}$ = 0.5 | 0.48 | | -0.004 | 0.02 | $F_{1, 8.12}$ = 0.03 | 0.87 | |
| | **Rainfall: age** | **-0.02** | **0.009** | **$F_{1, 1084.38}$ = 4.7** | **0.03** | | *-0.007* | *0.01* | *$F_{1, 457.48}$ = 0.16* | *0.69* | *1* |
| | *Rainfall: sex* | *0.009* | *0.009* | *$F_{1, 1067.76}$ = 1.1* | *0.30* | *1* | *-0.01* | *0.009* | *$F_{1, 458.07}$ = 2.3* | *0.13* | *2* |
| Yellow Brightness | | | | *City* | | | | | *Forest* | | |
| | *Intercept* | *73.2* | *52* | *$F_{1, 8.01}$ = 1.9* | *0.19* | | *66.8* | *54.7* | *$F_{1, 8.13}$ = 1.5* | *0.26* | |
| | **Sex** | **1.85** | **0.17** | **$F_{1, 719.95}$ = 121.7** | **<0.001** | | **21.1** | **9.84** | **$F_{1, 478.48}$ = 4.5** | **0.03** | |
| | Age | 0.008 | 0.17 | $F_{1, 1087.63}$ = 0.002 | 0.96 | | 0.26 | 0.23 | $F_{1, 468.72}$ = 1.2 | 0.28 | |
| | Temperature | -2.5 | 2.26 | $F_{1, 8.01}$ = 1.2 | 0.31 | | -2.28 | 2.53 | $F_{1, 8.13}$ = 0.8 | 0.39 | |
| | Temperature: sex | *0.04* | *0.34* | *$F_{1, 1066.50}$ = 0.02* | *0.89* | *1* | **-0.94** | **0.45** | **$F_{1, 477.96}$ = 4.3** | **0.04** | |
| | *Temperature: age* | *0.41* | *0.34* | *$F_{1, 1069.45}$ = 1.5* | *0.22* | *2* | *-0.71* | *0.49* | *$F_{1, 475.30}$ = 2.1* | *0.15* | *1* |
| | *Intercept* | *17.7* | *2.2* | *$F_{1, 8.19}$ = 66.5* | *<0.001* | | *0.69* | *0.05* | *$F_{1, 8.20}$ = 43.2* | *<0.001* | |
| | **Sex** | **2.09** | **0.31** | **$F_{1, 1080.23}$ = 45.6** | **<0.001** | | **0.65** | **0.24** | **$F_{1, 319.94}$ = 7.3** | **0.007** | |
| | Age | 0.006 | 0.17 | $F_{1, 1086.64}$ = 0.001 | 0.97 | | -0.64 | 0.48 | $F_{1, 480.71}$ = 1.7 | 0.19 | |
| | Rainfall | -0.70 | 1.25 | $F_{1, 8.07}$ = 0.3 | 0.59 | | -0.44 | 1.23 | $F_{1, 8.32}$ = 0.1 | 0.72 | |
| | Rainfall: sex | -0.16 | 0.17 | $F_{1, 1051.80}$ = 0.9 | 0.35 | | *0.22* | *0.22* | *$F_{1, 463.68}$ = 1.0* | *0.31* | *1* |

| | | | | | | | | | | | |
|---|---|---|---|---|---|---|---|---|---|---|---|
| | **Rainfall: age** | *0.07* | *0.18* | *$F_{1, 1082.55}$ = 0.1* | *0.70* | *1* | **0.51** | **0.24** | **$F_{1, 487.72}$ = 4.5** | **0.03** | |
| UV Chroma | *City* | | | | | | *Forest* | | | | |
| | *Intercept* | *0.19* | *0.2* | $F_{1, 8.02}$ = *0.9* | *0.37* | | *0.11* | *0.19* | $F_{1, 8.02}$ = *0.4* | *0.56* | |
| | **Sex** | **0.008** | **0.0009** | **$F_{1, 726.59}$ = 80.8** | **<0.001** | | **0.007** | **0.001** | **$F_{1, 322.27}$ = 39.1** | **<0.001** | |
| | **Age** | **0.003** | **0.0009** | **$F_{1, 1085.01}$ = 14.2** | **<0.001** | | **0.002** | **0.001** | **$F_{1, 464.52}$ = 5.1** | **0.02** | |
| | Temperature | -0.0005 | 0.009 | $F_{1, 8.02}$ = 0.003 | 0.96 | | 0.004 | 0.009 | $F_{1, 8.02}$ = 0.2 | 0.64 | |
| | *Temperature: sex* | *-0.002* | *0.002* | $F_{1, 1068.96}$ = *0.38* | *0.54* | *1* | *-0.002* | *0.002* | $F_{1, 483.80}$ = *0.8* | *0.36* | *1* |
| | *Temperature: age* | *-0.003* | *0.002* | $F_{1, 1065.41}$ = *2.1* | *0.15* | *2* | *-0.002* | *0.002* | $F_{1, 482.82}$ = *1.2* | *0.27* | *2* |
| | *Intercept* | *0.18* | *0.008* | $F_{1, 8.76}$ = *506.2* | *<0.001* | | *0.21* | *0.009* | $F_{1, 8.25}$ = *608.9* | *<0.001* | |
| | **Sex** | **0.008** | **0.0009** | **$F_{1, 726.64}$ = 80.8** | **<0.001** | | 0.002 | 0.002 | $F_{1, 491.01}$ = 1.3 | 0.26 | |
| | **Age** | **0.003** | **0.0009** | **$F_{1, 1085.04}$ = 14.2** | **<0.001** | | **0.002** | **0.001** | **$F_{1, 461.47}$ = 5.3** | **0.02** | |
| | Rainfall | -0.002 | 0.005 | $F_{1, 8.00}$ = 0.2 | 0.67 | | -0.005 | 0.004 | $F_{1, 8.30}$ = 1.7 | 0.23 | |
| | **Rainfall: sex** | *0.0006* | *0.0009* | $F_{1, 1042.19}$ = *0.5* | *0.47* | *2* | **0.002** | **0.0009** | **$F_{1, 457.19}$ = 6.4** | **0.01** | |
| | *Rainfall: age* | *-0.0002* | *0.0009* | $F_{1, 1080.08}$ = *0.04* | *0.84* | *1* | *0.0001* | *0.001* | *$F_{1, 485.97}$ = 0.02* | *0.89* | *1* |

**Table S5**: Table of the association between linear climatic variables and colouration traits in forest and urban great tits, separated per sex or age, depending on the significant interaction of the models when population were separated (Table S4). Type III sums of squares Anova are used and P-values are obtained with F-tests. Significant fixed effects are in bold. Reference levels of sex and age are females and yearlings.

| | | | | *ANOVA (main effects)* | |
|---|---|---|---|---|---|
| Response variable | Fixed effect | Estimates | *SE* | *F* | *Pr (>F)* |
| *Chroma: females (city) N=568* | *Intercept* | *1.29* | *2.2* | $F_{1, 8.03} = 0.36$ | *0.56* |
| | **Age** | **0.02** | **0.01** | $\mathbf{F_{1, 554.97} = 4.7}$ | **0.03** |
| | Temperature | -0.03 | 0.09 | $F_{1, 8.03} = 0.09$ | 0.76 |
| *Chroma: males (city) N=534* | *Intercept* | *1.98* | *2.01* | $F_{1, 8.03} = 0.97$ | *0.35* |
| | **Age** | **0.04** | **0.01** | $\mathbf{F_{1, 520.50} = 9.6}$ | **0.002** |
| | Temperature | -0.06 | 0.09 | $F_{1, 8.03} = 0.5$ | 0.50 |
| *Chroma: adults (city) N=693* | *Intercept* | *0.63* | *0.08* | $F_{1, 8.03} = 57.9$ | *<0.001* |
| | **Sex** | **-0.03** | **0.01** | $\mathbf{F_{1, 409.57} = 5.9}$ | **0.02** |
| | Rainfall | 0.01 | 0.05 | $F_{1, 7.99} = 0.1$ | 0.76 |
| *Chroma: yearlings (city) N=409* | *Intercept* | *0.57* | *0008* | $F_{1, 8.52} = 48.5$ | *<0.001* |
| | **Sex** | **-0.05** | **0.02** | $\mathbf{F_{1, 395.61} = 9.6}$ | **0.002** |
| | Rainfall | 0.04 | 0.05 | $F_{1, 8.01} = 0.6$ | 0.48 |
| *Brightness: females (forest) N=267* | *Intercept* | *65.3* | *51.4* | $F_{1, 8.02} = 1.6$ | *0.34* |
| | Age(adults) | 0.48 | 0.30 | $F_{1, 256.759} = 2.5$ | 0.12 |
| | Temperature | -2.00 | 2.39 | $F_{1, 8.02} = 0.9$ | 0.38 |
| *Brightness: males (forest) N=237* | *Intercept* | *89.4* | *57.9* | $F_{1, 8.03} = 2.4$ | *0.16* |
| | Age | -0.07 | 0.36 | $F_{1, 214.7} = 0.04$ | 0.84 |
| | Temperature | -3.01 | 2.70 | $F_{1, 8.02} = 1.5$ | 0.25 |
| *Brightness: adults (forest) N=275* | *Intercept* | *17.9* | *2.9* | $F_{1, 8.061} = 38.4$ | *<0.001* |
| | Sex(males) | 0.27 | 0.32 | $F_{1, 180.092} = 0.7$ | 0.40 |
| | Rainfall | 0.05 | 1.28 | $F_{1, 7.986} = 0.001$ | 0.97 |
| *Brightness: yearlings (forest) N=229* | *Intercept* | *18.0* | *2.7* | $F_{1, 8.077} = 45.0$ | *<0.001* |
| | **Sex** | **1.03** | **0.34** | $\mathbf{F_{1, 218.198} = 9.0}$ | **0.003** |
| | Rainfall | -0.39 | 1.19 | $F_{1, 8.204} = 0.1$ | 0.75 |
| *UV Chroma: females (forest) N=267* | *Intercept* | *0.21* | *0.09* | $F_{1, 8.073} = 534.4$ | *<0.001* |
| | Age | 0.003 | 0.001 | $F_{1, 225.961} = 3.5$ | 0.06. |
| | Rainfall | -0.005 | 0.004 | $F_{1, 8.095} = 1.6$ | 0.24 |
| *UV Chroma: males (forest) N=237* | *Intercept* | *0.21* | *0.008* | $F_{1, 8.134} = 714.2$ | *<0.001* |
| | Age | 0.002 | 0.001 | $F_{1, 224.170} = 1.9$ | 0.17 |
| | Rainfall | -0.002 | 0.004 | $F_{1, 8.047} = 0.5$ | 0.21 |

**Table S6**: Table of the association between: a/ proportion of newly ringed birds (i.e., immigrants) and years in each habitat; b/ age structure (approximated by the percentage of yearlings) and years in each habitat. Type III sums of squares Anova are used and P-values are obtained with F-tests. Reference levels of habitats is the city

| | | | | *ANOVA (main effects)* | |
|---|---|---|---|---|---|
| Response variable | Fixed effect | Estimates | *SE* | *F* | *Pr (>F)* |
| *a/ Percentage of newly ringed birds* | *Intercept* | *56.2* | *4.2* | $F_{1, 16} = 31615$ | *<0.001* |
| | Year | -8.81 | 4.3 | $F_{1, 16} = 737$ | 0.06 |
| | Habitat (forest) | -6.91 | 5.9 | $F_{1, 16} = 97.3$ | 0.26 |
| | Year: habitat | 4.53 | 6.1 | $F_{1, 16} = 97.3$ | 0.47 |
| *b/ Percentage of yearlings* | *Intercept* | *36.7* | *3.9* | $F_{1, 16} = 13472$ | *<0.001* |
| | Year | 2.9 | 3.9 | $F_{1, 16} = 80.2$ | 0.47 |
| | Habitat (forest) | 6.9 | 5.5 | $F_{1, 16} = 241$ | 0.22 |
| | Year: habitat | -1.5 | 5.6 | $F_{1, 16} = 10.1$ | 0.79 |

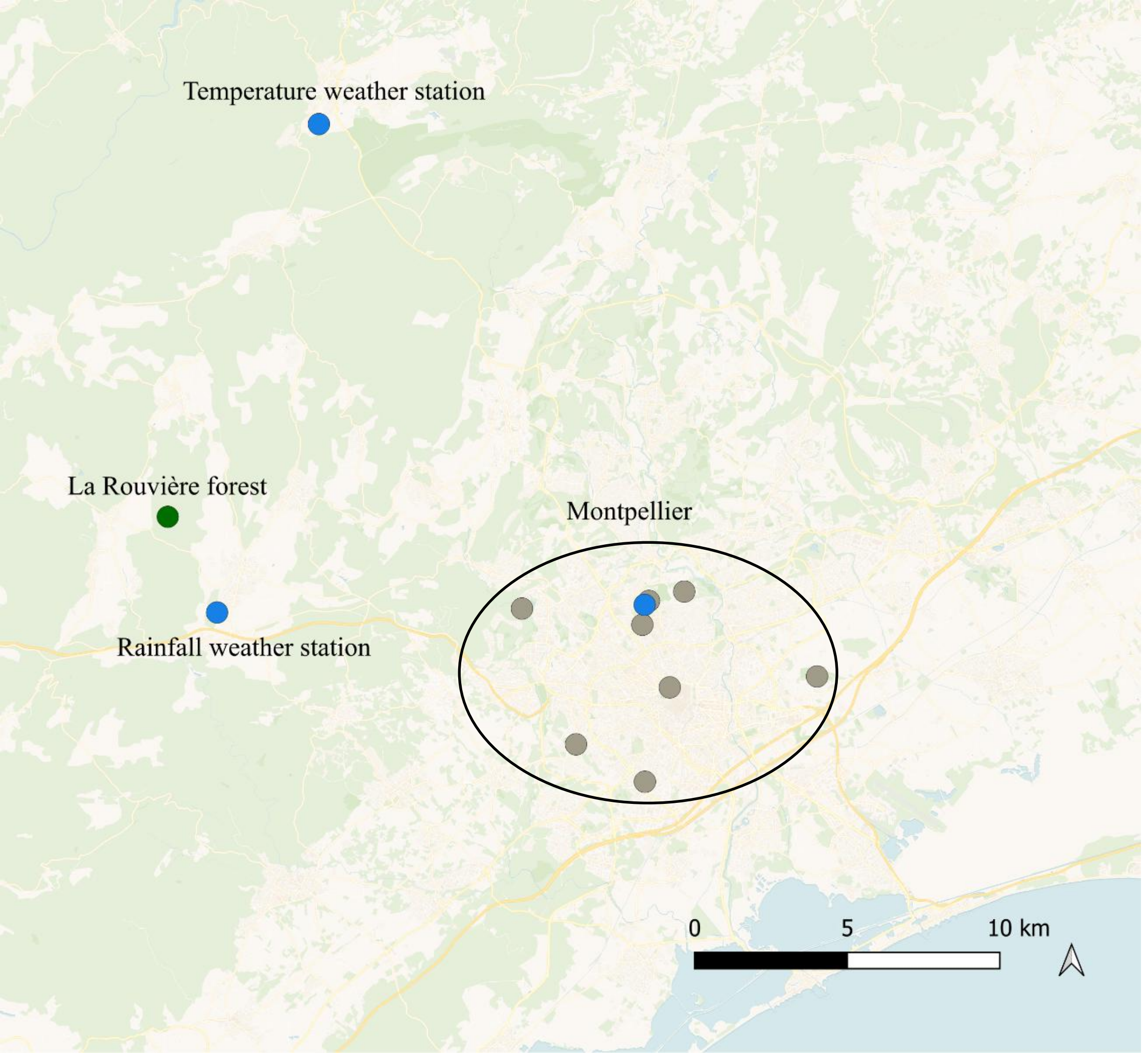


**Fig.S1**: Map of the Montpellier area (southern France) showing the urban and forest great tit populations and the meteorological stations used in the study. The eight urban sampling sites are shown as grey dots within the black ellipse surrounding the city of Montpellier. The forest population of La Rouvière is indicated by the green dot. Blue dots represent the meteorological stations from which climatic data were obtained: the "Rainfall weather station" (Montarnaud) and the "Temperature weather station" (Saint-Martin-de-Londres) located near the forest site, and the "Montpellier weather station" located within the urban area.

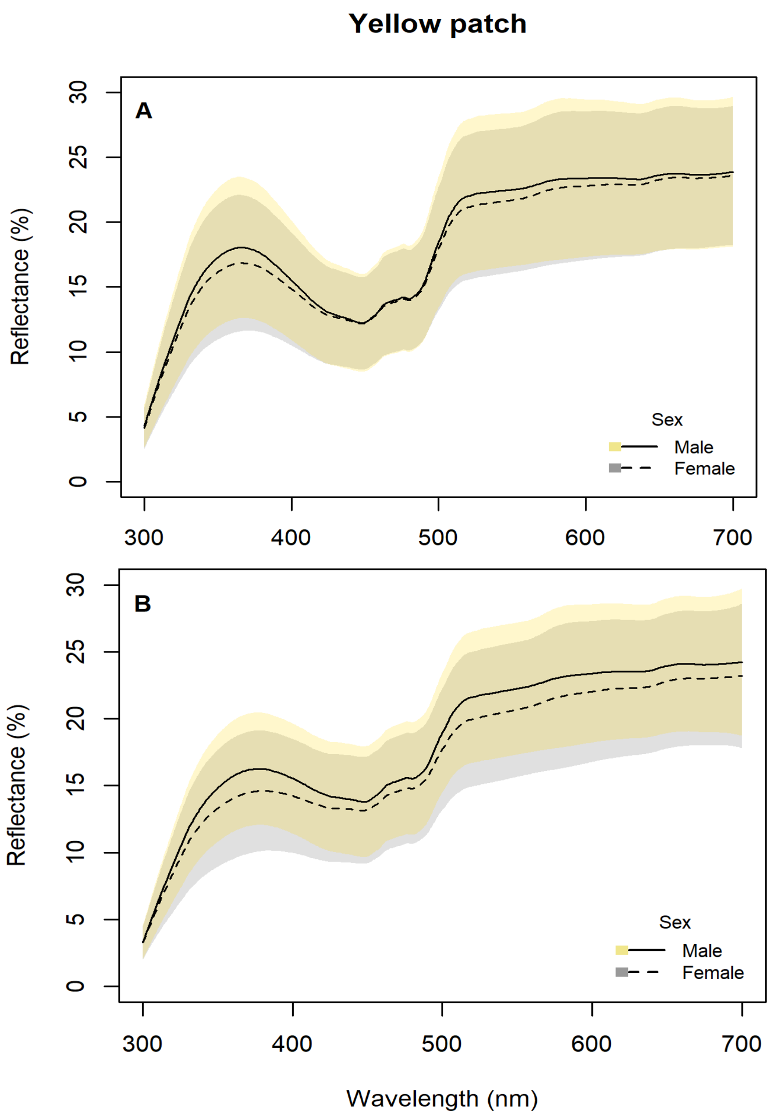


**Fig.S2**: Average spectra obtained by spectrophotometry of 4 feathers from the yellow breast of forest (A) and city (B) great tits. Lines represent the mean spectra for both sexes. The range of spectra obtained are represented as coloured stripes for both sexes.

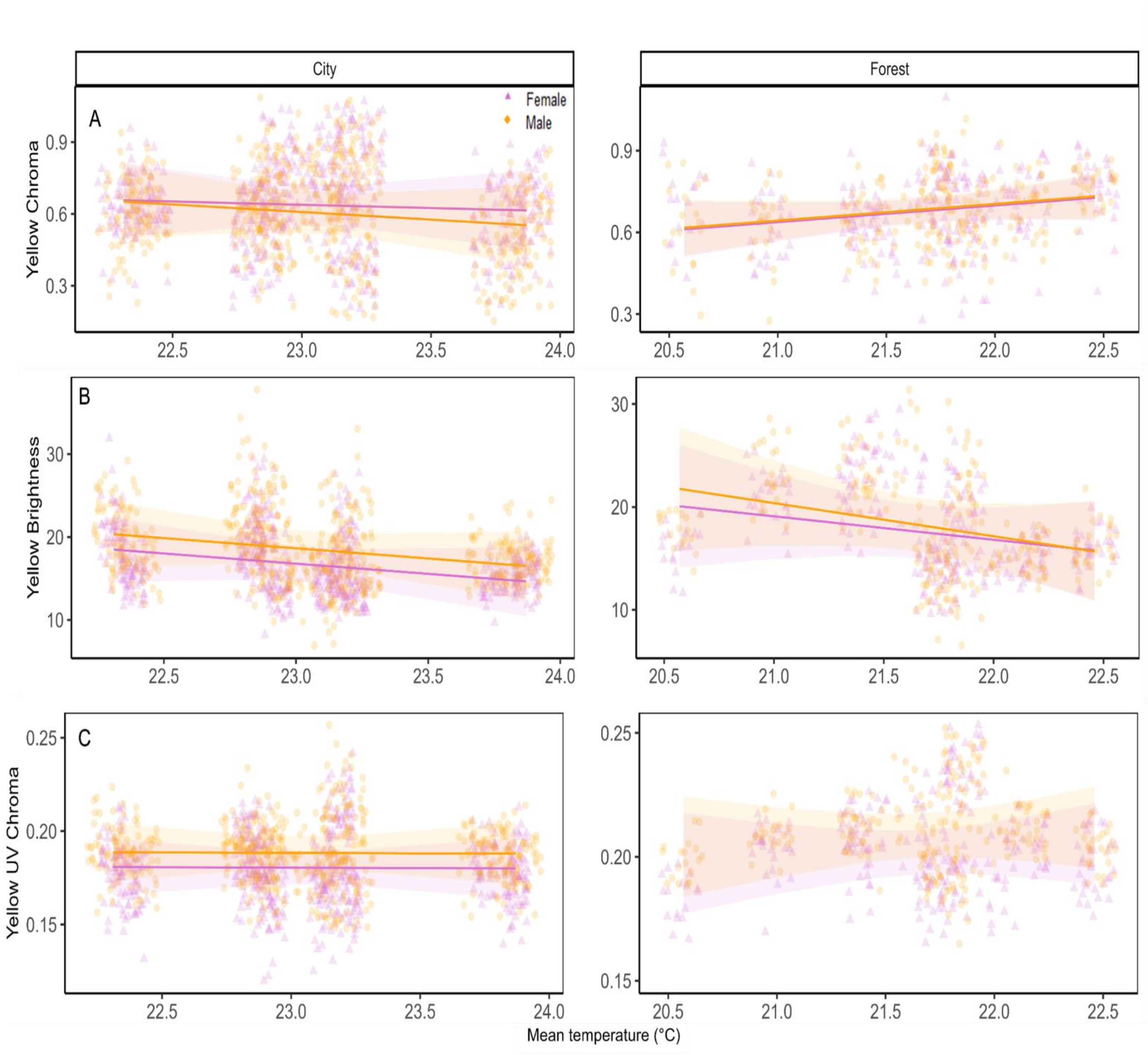


**Figure S3:** Association of male and female great tits yellow breast colouration of the forest and the city with mean temperature (°C) during the moulting period (Table S6), (A) Yellow chroma, (B) Brightness and (C) UV Chroma. The lines show the predicted slope values. The dots show the raw data.95% confidence intervals are shown as the coloured stripes. A jitter was added to visualise overlapping points. Violet triangles and dashed lines represent the females, orange dots and solid lines are for the males. Red squares and dashed lines represent the yearlings, blue diamonds and solid lines are for adults.

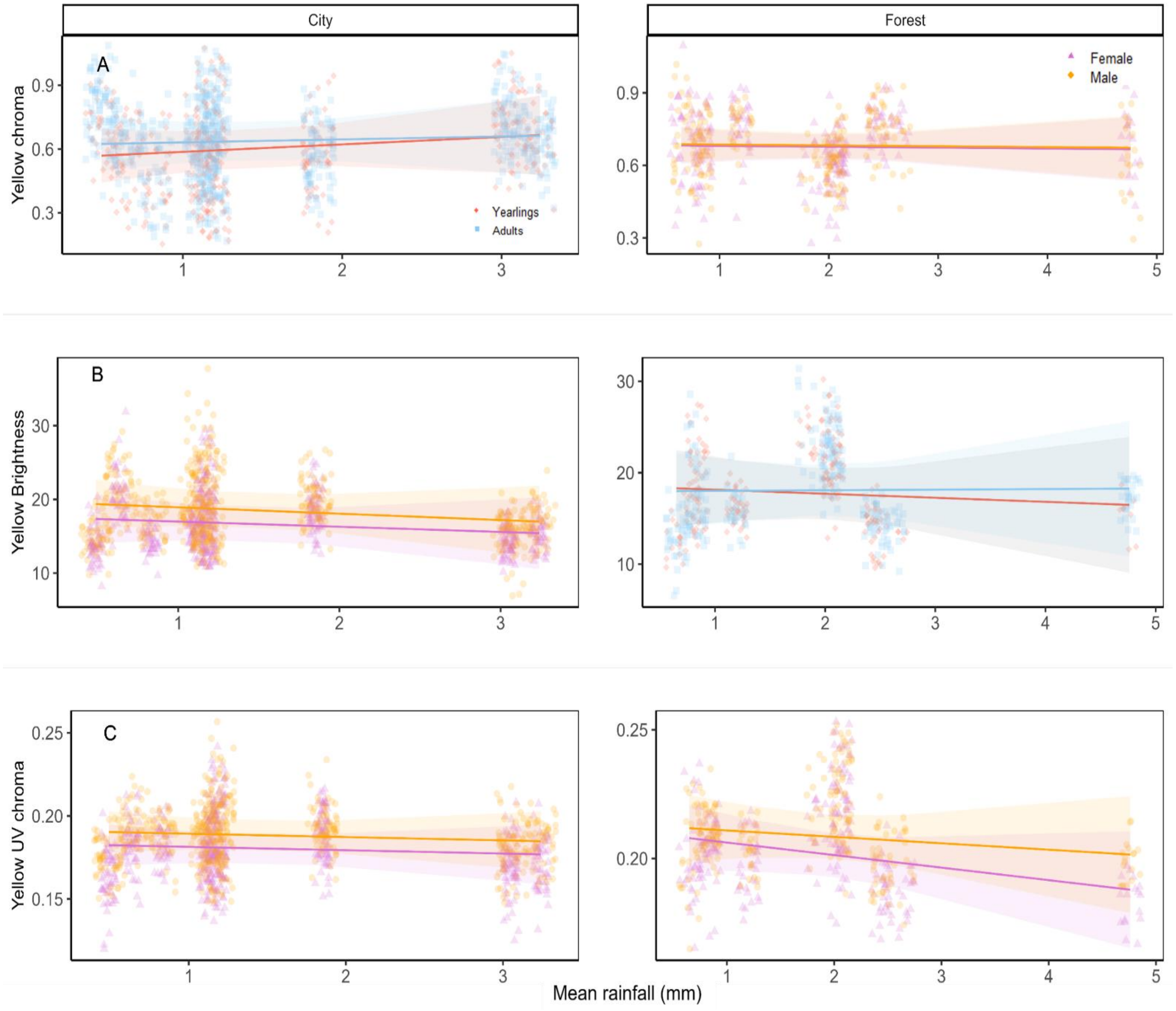


**Figure S4:** Association of male and female great tits yellow breast colouration of the forest and the city with mean rainfall (mm) during the moulting period (Table S6), (A) Yellow chroma, (B) Brightness and (C) UV Chroma. The lines show the predicted slope values. The dots show the raw data.95% confidence intervals are shown as the coloured stripes. A jitter was added to visualise overlapping points. Violet triangles and dashed lines represent the females, orange dots and solid lines are for the males. Red squares and dashed lines represent the yearlings, blue diamonds and solid lines are for adults.

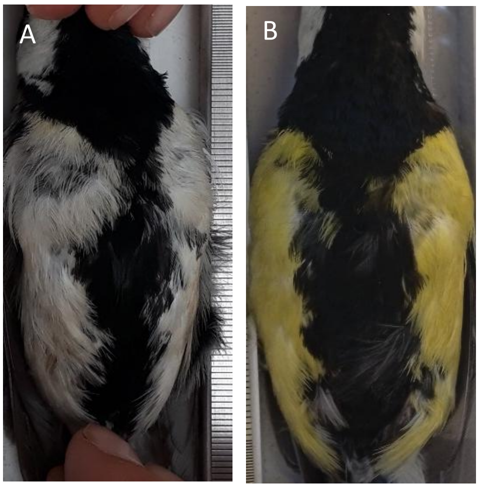


**Figure S5**: On the right, pictures of urban (A) and rural (B) male great tits yellow breasts from our populations

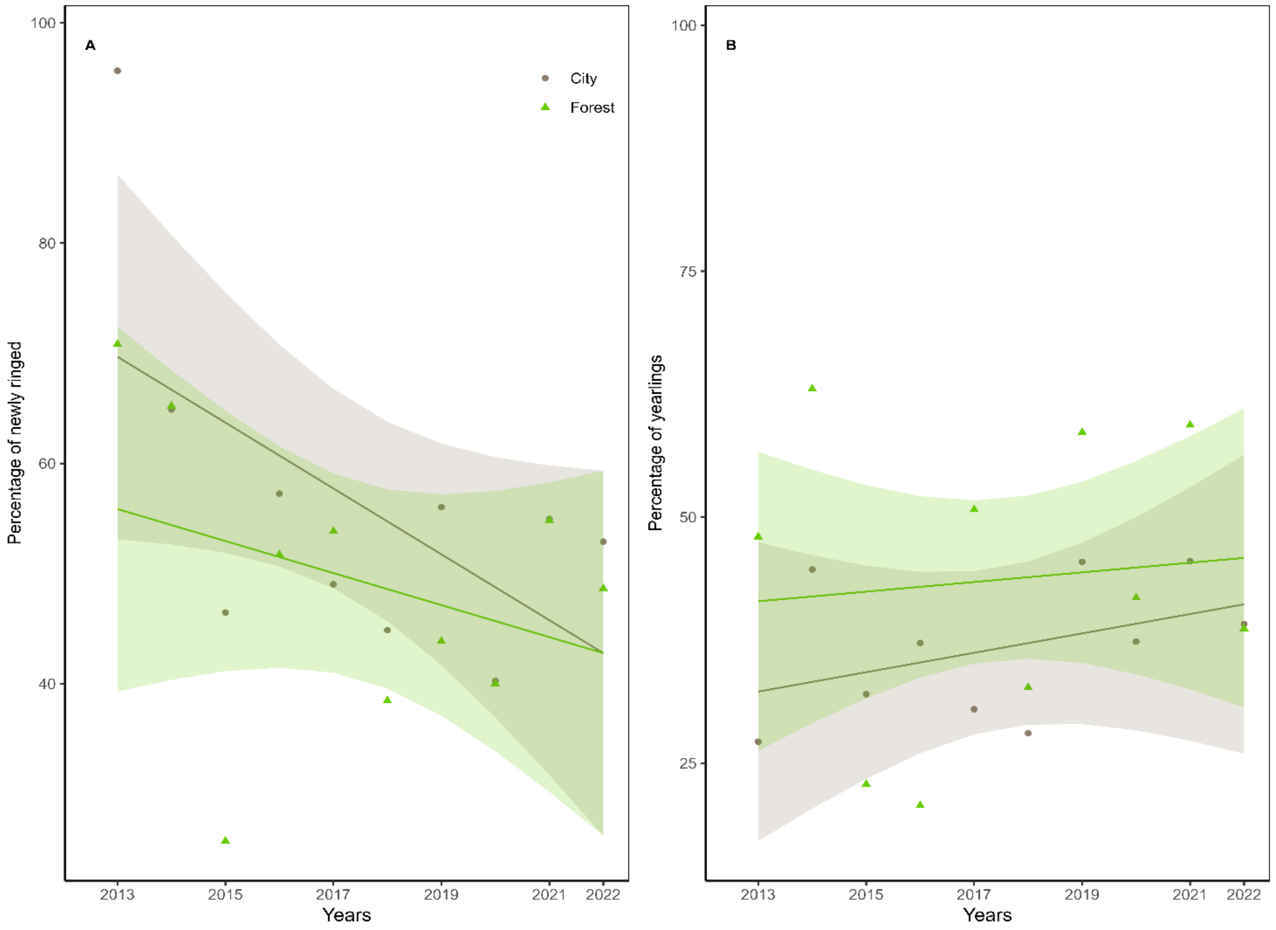


**Figure S6:** Temporal variation of the percentage of immigrants (newly ringed, A) and yearlings (B) in both habitats. The lines show the predicted slope values. The dots show the raw data.95% confidence intervals are shown as the coloured stripes. Green triangles and lines represent the forest, grey dots and lines are for the city
